# Plasma protein biomarkers to detect early gastric preneoplasia and cancer: a prospective study

**DOI:** 10.1101/2025.04.29.651246

**Authors:** Quentin Giai Gianetto, Valérie Michel, Thibaut Douché, Karine Nozeret, Aziz Zaanan, Oriane Colussi, Isabelle Trouilloud, Simon Pernot, Marie-Noelle Ungeheuer, Catherine Julié, Nathalie Jolly, Julien Taïeb, Dominique Lamarque, Mariette Matondo, Eliette Touati

## Abstract

Gastric cancer (GC) often presents a poor prognosis due to its asymptomatic phenotype at early stages. Upper endoscopy, the current gold standard for diagnosis, is invasive with limited sensitivity for detecting gastric preneoplasia. Non-invasive biomarkers, such as blood circulating proteins offer a promising alternative for an early detection. Using mass spectrometry-based proteomics we identified plasma proteins as biomarkers of the presence of gastric preneoplasia and cancer lesions in an exploratory subgroup of patients (n=39). Fifteen promising protein candidates emerged to distinguish patient categories and were further confirmed by enzyme-linked immunosorbent assays (ELISA) in plasma samples from a cohort of 138 participants. Our predictive models demonstrated high classification performance with a minimal set of biomarkers, making them clinically applicable. Repeated cross-validations yielded high Area Under the Receiver Operating Characteristics (AUROC) values, notably distinguishing cancerous or precancerous cases from non-cancerous ones. Leveraging simple blood sampling, this strategy holds promise to detect high-risk gastric lesions, even at asymptomatic stages. Such an approach could significantly improve early detection and clinical management of GC, offering direct benefit for patients.

## Introduction

Gastric cancer (GC) is the fifth most commonly diagnosed cancer and the fourth leading cause of cancer-related death worldwide (Bray *et al*, 2018). With over 1 million new cases reported annually, GC remains an important healthcare challenge. It predominantly affects individuals over the age of 60, preferentially men, and it is strongly associated with *H. pylori* infection (Mukaisho *et al*, 2015). *H. pylori* infects nearly half of the global population and is recognized as a major risk factor for GC (Moss, 2017). While the global incidence and mortality of GC have declined over the past decades, a concerning rise in cases among younger individuals (< 50 years) has been observed in both low- and high-income countries (Arnold *et al*, 2020) (Anderson *et al*, 2018). Due to its asymptomatic phenotype during the first steps of its development, GC is mainly associated with a poor prognosis, highlighting the importance of its early detection. Effective early detection strategies are particularly critical with important benefits for patient outcomes, as observed in countries with established screening programs (Sun *et al*, 2024). The incidence of GC is higher in Eastern Asia, as in Japan and South Korea where national screening programs have been implemented leading to a 5-year survival of about 60%, compared to 15-20% in Western countries, thus highlighting the benefit of prevention (Shen *et al*, 2013).

The majority of GC cases are adenocarcinoma that develop through a multistep process, initiated by a chronic inflammation and progressing through non-atrophic gastritis (NAG) also referred to gastritis, atrophic gastritis (AG), intestinal metaplasia (IM) and dysplasia (DYS) preceding cancer lesions (Correa, 1992). If detected at an early stage, meaning AG and/or low-grade preneoplasia, GC can be a preventable disease (Rugge *et al*, 2017), making the identification of these early-stage lesions a key clinical priority. The prevalence of *H. pylori*- associated AG and IM, increases logarithmically with age and confers higher risk of GC (Graham & Lee, 2025). While *H. pylori* eradication can reduce the risk of GC, residual risk persists, especially in individuals with existing gastric preneoplasia or high baseline risk (Liou *et al*, 2024) (Ford *et al*, 2022). Presently, upper endoscopy is the gold standard for GC diagnosis and preneoplasia monitoring (Pimentel-Nunes *et al*, 2019) (Dinis-Ribeiro *et al*, 2024). However, this invasive method is costly and often limited in detecting transient lesions such as DYS (Pimentel-Nunes *et al*, 2019) (Dinis-Ribeiro *et al*, 2024). It has been reported that 10-20% of patients with GC had no preneoplastic lesions detected by endoscopy within 6 to 36 months prior to the diagnosis (Beck *et al*, 2021), underscoring the limitations of current diagnostic approaches and the urgent need for complementary detection methods. Therefore, the discovery of liquid biopsy-based biomarkers as in blood is of paramount interest to identify the presence of gastric lesions at an asymptomatic stage. There is a real need in clinic, not only for screening and early detection of GC but also for patient follow-up and treatment monitoring.

Circulating proteins have emerged as promising biomarkers for cancer detection, as previously reported for the CancerSeek test (Cohen *et al*, 2018). Although glycoproteins including carbohydrate antigen 12-5 (CA12-5), 19-9 (CA19-9) and carcinoembryonic antigen (CEA) are routinely used in clinical practice, their sensitivity and specificity for GC remain limited (Desai *et al*, 2023). Particularly for early GC, where these conventional markers show extremely low positive rates – ranging from merely 2.5% to 15.4% of cases associated with elevated marker levels (Feng *et al*, 2017). Serological markers like the well-studied pepsinogen I and II (PGI and PGII) and PGI/PGII ratio show some predictive value for corpus AG with a sensitivity from 67% to 85% and a specificity from 73% to 87% (Agreus *et al*, 2012) (Loong *et al*, 2017). Despite many efforts, none of the reported blood-based GC biomarkers have been translated into clinical practice to date. This is mainly due to their insufficient sensitivity for early-stage lesions, the lack of large-scale validation studies, and the challenges in standardization across different patient populations.

Recent improvements in high-throughput technologies, particularly mass spectrometry-based proteomics, have enabled the identification of biomarker panels with higher diagnostic accuracy compared to single proteins. For instance, a signature of 19 proteins has been identified to distinguish early-stage (I/II) GC from healthy subjects (Shen *et al*, 2019). However, until now, there is a lack of studies that have specifically investigated blood biomarkers across the full spectrum of lesions that composed the gastric carcinogenesis cascade, from NAG through preneoplasia (AG, IM and DYS) to cancer (Bazin *et al*, 2024).

Our study addresses this gap, using liquid chromatography coupled to tandem mass spectrometry (LC-MS/MS) proteomics to identify plasma protein signatures associated with each stage of gastric lesions progression. We specifically focused on identifying biomarkers capable of detecting preneoplastic lesions, a key point where treatment could prevent progression to malignancy. After the discovery of some proteins by LC-MS/MS, we validated promising biomarker candidates using ELISA in a larger cohort of patients, enabling us to develop and test predictive models with potential clinical applicability. This comprehensive approach offers several advantages over previous studies: it examines multiple disease stages rather than binary comparisons, employs rigorous validation in a substantial patient cohort and prioritizes biomarkers with practical clinical utility. Our findings provide novel insights into protein biomarker panels that could significantly improve risk stratification and early detection for individuals at risk of GC, potentially transforming the clinical management of this deadly disease.

## Results

### Characteristics of the studied cohort and patient diagnosis

The general characteristics of the patients included in the cohort are reported in Table 1. All patients were previously diagnosed with upper gastrointestinal (GI) endoscopy, and those with AG had undergone upper GI endoscopy within 12 months prior to inclusion. Endoscopic examination of the mucosa was performed under white and blue light, in order to obtain biopsies from sites with the most severe lesions of atrophy or metaplasia, in accordance with European recommendations (Pimentel-Nunes *et al*, 2019). Eighteen biopsy samples were systematically taken from the four sides of the antrum, angulus and the gastric body. The group of preneoplasia is heterogenous and includes patients with AG (21%), IM (47%) and both IM and DYS (32%). 58% and 61% of patients with NAG or preneoplasia respectively, were *H. pylori*-positive and 28% in the GC group. All healthy individuals in the control group are *H. pylori*-negative with mean age significantly lower compared to gastritis, preneoplasia and GC patients (Table 1).

**Table 1:** Characteristics of the study population. The gastritis group corresponds to patients with non-atrophic gastritis. The group of preneoplasia is composed of AG (21%), IM (47%) and IM and DYS (32%). GC group includes GC of intestinal type (53%), of diffuse type (37%) and undifferentiated (10%), among which stage II (2%), III (29%) and IV (69%). The exploratory LC-MS/MS analysis was performed on a subgroup of patients (numbers in italic). *P* values for statistical analysis using Mann-Whitney test to compare age in healthy groups to other patient groups.

| Group | n | Mean age (range) | Sex ratio (M/F) | *H. pylori* positive (%) |
| --- | --- | --- | --- | --- |
| Healthy | 32 | 42 (21-70) | 0.4 | 0 |
|  | *10* | *38 (22-69)* | *0.4* | *0* |
| Gastritis | 26 | 59 (27-88)<br>*P=0.0003* | 0.7 | 58% |
|  | *10* | *38 (22-69)*<br>*P=0.0015* | *0.6* | *58%* |
| Preneoplasia | 20 | 70 (50-83)<br>*P<0.0001* | 0.4 | 61% |
|  | *9* | *70 (54-83)*<br>*P=0.0006* | *0.5* | *50%* |
| Gastric cancer | 60 | 62 (29-84)<br>*P<0.0001* | 2 | 28% |
|  | *10* | *62 (51-74)*<br>*P=0.0044* | *4* | *40%* |
| Total | **138** |  |  |  |
|  | **39** |  |  |  |

#### Multivariate analysis of the LC-MS/MS-based proteomics data

As illustrated in Figure 1, a first exploratory phase was conducted using LC-MS/MS-based proteomics on plasma from 39 individuals of the cohort, including healthy and patients diagnosed with gastritis, preneoplasia and GC (Table 1). Following plasma protein preparation (see Supplementary material), a total of 691 proteins were identified and quantified.

**Figure 1:**
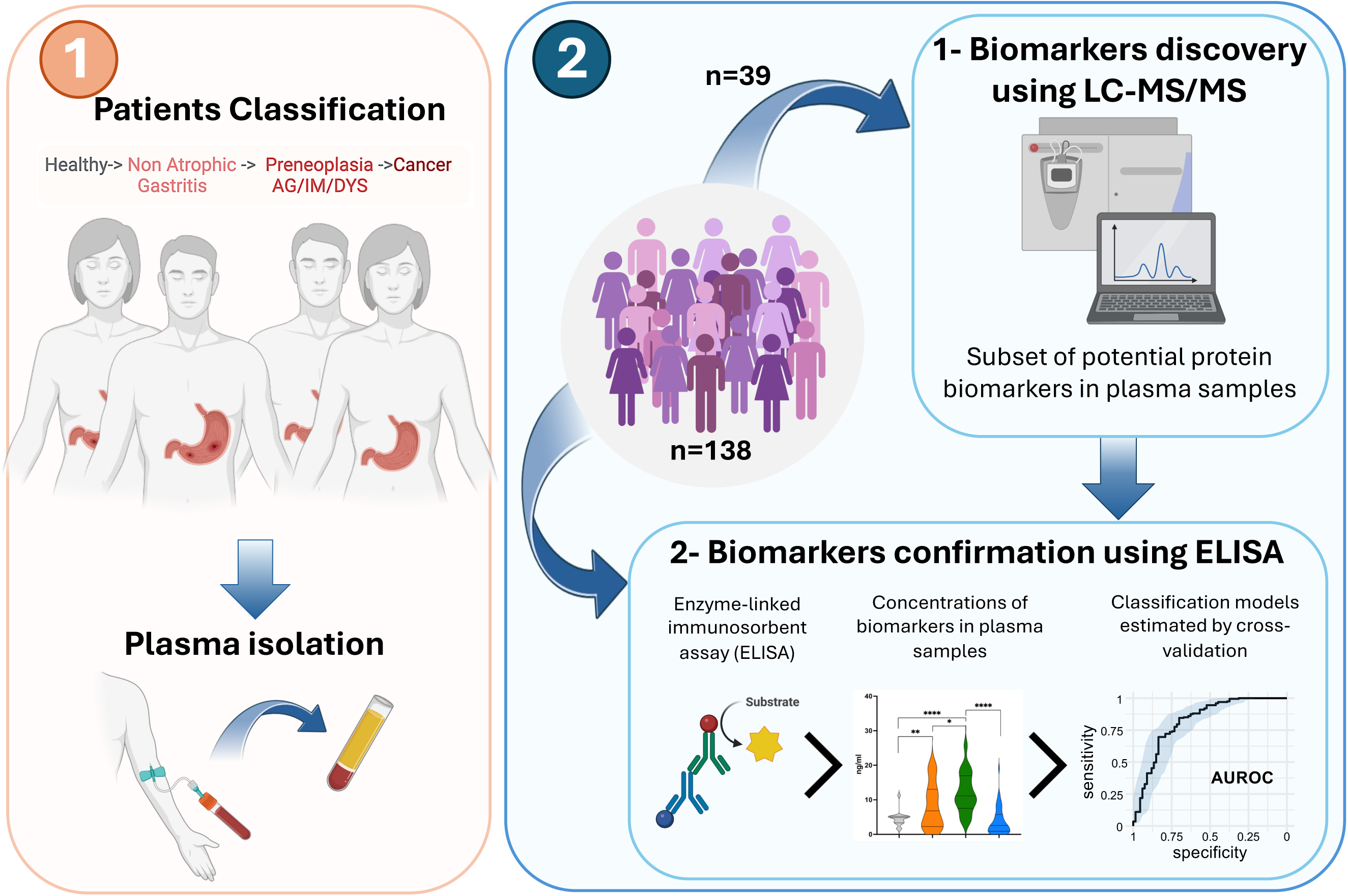
Schematic representation of the study worflow. Overview of 1) the setting up of a new multicenter cohort (n=138) including a group of healthy individuals and 3 groups of patients with gastritis, preneoplasia and GC. The group of patients with preneoplasia corresponds to AG either with IM or DYS or not. From each patient blood sample is collected and plasma isolated, and of 2) the exploratory proteomic analysis by LC-MS/MS on plasma samples from a subgroup (n=39) of patients and healthy subjects and their confirmation by ELISA through mAUROC analysis and prediction models validation.

To assess the similarity of proteomic profiles across groups, correlation analysis was performed (Figure 2). Overall, the proteomes showed high levels of correlation, with a minimum of 80.05%. Within-group correlation was particularly strong, especially among patients in the same diagnostic category. Notably, proteomes from gastritis and preneoplasia patients were closely aligned, whereas those from healthy individuals and GC patients are more distinct from other groups of pathology. Partial Least Square-Discriminated Analysis (PLS-DA) was used to further explore group separation based on proteomic profiles. The first two PLS-DA components clearly distinguished healthy individuals from patients affected by a pathology. Components 2 and 3 allow distinguishing GC patients from others (Figure 2A). As observed from the correlation analysis (Figure 2B), the separation between patients with gastritis from those with preneoplasia is low but still occurs at the level of the fourth component of the PLS-DA that explains only 3% of the total variance (Figures 2A). Together, the first four PLS-DA components allowed for the discrimination of the different patient categories based on their proteomes. These findings indicate distinct proteomic signatures in plasma samples associated with each group of patients.

**Figure 2.**
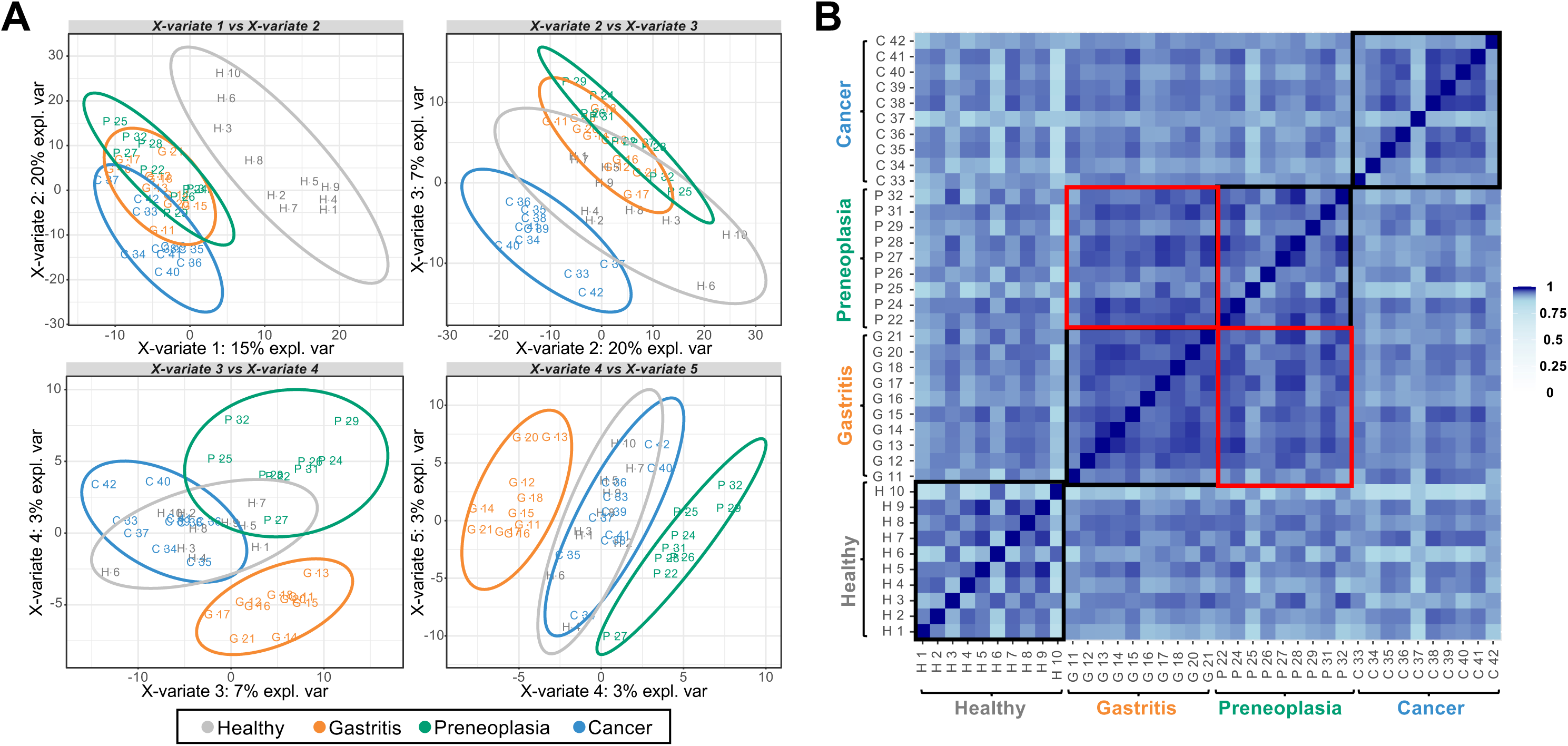
Multivariate analysis of the mass spectrometry-based proteomics data. A Sample plot of the PLS-DA model using 5 components. These components explain 48% of the total inertia. B Pairwise correlations between samples. Shades of blue represent correlation values. Black squares are sub-matrices corresponding to each category of patients. Red squares are the correlation matrices between samples of patients affected by gastritis and preneoplasia (AG/P).

#### Biomarker candidates identified by LC-MS/MS-based data

To identify candidate biomarkers capable of distinguishing between patient groups, we performed differential analyses using multiple moderated t-tests for all pairwise comparisons (“*Two by two”* comparisons), as well as a sparse PLS-DA analysis, as described in the Methods section.

##### “Two by two” comparisons

For comparison of one pathology *vs* another, two types of statistical analyses were conducted: a “relaxed” analysis by imposing for each protein, at least two quantified values in the plasma samples of patients from one of the two compared categories (Figure 3A), and a “strict” analysis by imposing at least nine quantified values in one of the two compared categories (Figure 3B). Since these proteins are quantified in almost all patients in a category, we can expect the “strict” analysis to give more robust results. Six different comparisons were performed: each group of patients *vs* healthy, gastritis *vs* preneoplasia, and gastritis or preneoplasia *vs* GC. From the pairwise comparisons resulting from the “relaxed” analysis (Figure 3A), 408 proteins were found significantly more abundant in one category of patients than in another, among which 309 with differential level in at least two comparisons (see Supplementary Figure S1 and additional file 1). With the “strict” analysis approach (Figure 3B), 236 proteins show differential level comparing two groups: 149 are differential in at least two comparisons (see Supplementary Figure S2 and additional file 2). It should be noted that the comparison of preneoplasia and gastritis categories show the fewest proteins differentially abundant, with 58 differential proteins found in the “relaxed” analysis (Figure 3A, Supplementary Figure S1 and additional file 1), while none is found with the “strict” analysis (Figure 3B, Supplementary Figure S2 and additional file 2). Considering the “strict” analysis, 2 proteins appear with differential level in 5 comparisons over 6: Transferrin Receptor Protein 1 (TFRC) and Immunoglobulin heavy constant γ1 (IGHG1); 8 proteins in 4 comparisons over 6: Lysosomal Pro-X carboxypeptidase (PRCP), Serum amyloid A-1 protein (SAA1), Serum amyloid A-2 protein (SAA2), Serine protease 3 (PRSS3), α-mannosidase 2A1 (MAN2A1), Dermcidin (DCD), Glycosylphosphatidylinositol phospholipase D (GPLD1) and Interleukin-1 receptor accessory protein (IL1RAP). These proteins could serve as promising biomarker candidates for distinguishing between patient categories. In addition to pairwise condition comparisons, we performed a sparse PLS-DA analysis to identify a refined set of proteins capable of distinguishing all patient groups effectively.

**Figure 3.**
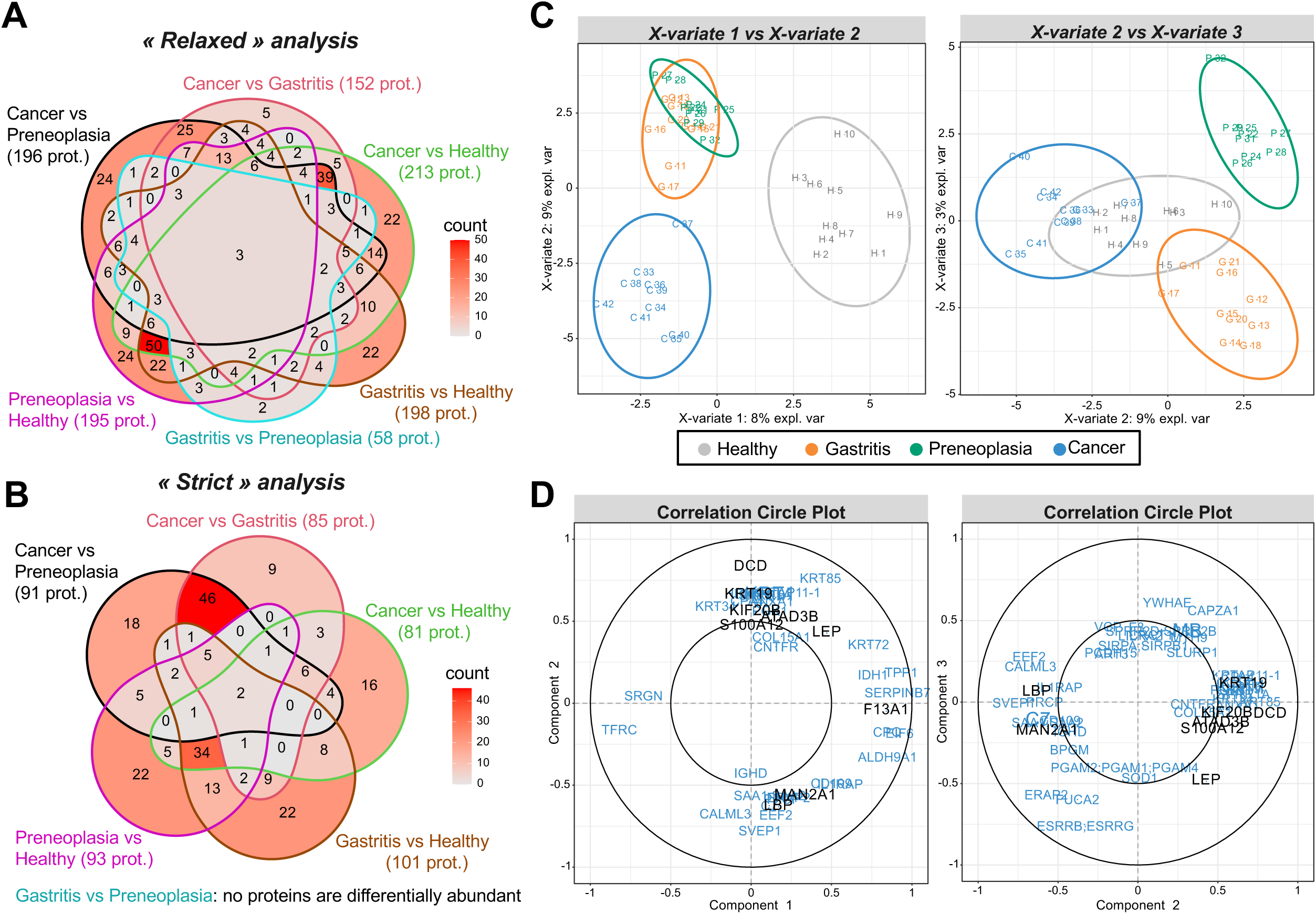
Identification of protein biomarkers from mass spectrometry-based proteomics data. A Venn diagrams summarizing the 408 biomarkers found in all the comparisons when imposing at least 2 quantified values in a category of patients (« relaxed » analysis): 309 are differential in at least two comparisons. B Venn diagrams summarizing the 236 biomarkers found in all the pairwise comparisons when imposing at least 9 quantified values in a category of patients (« strict » analysis): 149 are in at least two comparisons. C Sample plot of the sparse PLS-DA model with 3 components. These components explain 20% of the total inertia. Each plot represents the samples projected on two axes, each one corresponding to a latent component of the model. D Correlation circle plot of the sparse PLS-DA. The proteins measured by ELISA test are highlighted in black.

#### Sparse PLS-DA

Unlike the previous approach that compares each category of patients two by two, this sparse PLS-DA analysis attempts to separate all the groups according to a restricted number of proteins: we imposed the choice of only 10 biomarkers by component of the PLS-DA. The first component distinguishes healthy subjects from others, while the second one separates GC patients from those with gastritis and preneoplasia (Figure 3C). The third component allows a separation between gastritis and preneoplasia. The correlation circle plots highlight the correlation of each protein selected by the sparse PLS-DA with its different components (Figure 3D). Among the 85 proteins identified as potential biomarkers by the sparse PLS-DA analysis, Coagulation factor XIII A chain (F13A1) is highly correlated with the first component, while TFRC is inversely correlated. They could thus be biomarkers to separate healthy individuals from others. Other proteins like DCD, Keratin type I cytoskeletal 19 (KRT19), Kinesin-like protein KIF20B (KIF20B), ATPase family AAA domain containing 3B (ATAD3B), S100 calcium-binding protein A12 (S100A12) or Leptin (LEP) are positively correlated with the second component while MAN2A1, Lipopolysaccharide-binding protein (LBP) or Sushi Von Willebrand Factor Type A, EGF and Pentraxin Domain containing 1 (SVEP1) are inversely correlated. Thus, these proteins could contribute to separate GC patients from gastritis and preneoplasia patients. When examining proteins correlated with the third component separating gastritis and preneoplasia, Capping Actin Protein of Muscle Z-Line Subunit α (CAPZA1), α-L- Fucosidase 2 (FUCA2), Endoplasmic Reticulum Aminopeptidase 2 (ERAP2) and LEP are found. They are also positively or negatively correlated with the second component separating GC patients from others. Interestingly, LEP is the only protein kept by the sparse PLS-DA with an absolute correlation close or superior to 0.5 with the three components. It was also found differentially abundant in the “relaxed” analysis in four comparisons (GC *vs* gastritis, GC *vs* healthy, preneoplasia *vs* healthy and GC *vs* preneoplasia) (Supplementary material additional file 1) and could thus be an interesting candidate. Importantly, most of the proteins highlighted by the sparse PLS-DA, are categorized as potential prognostic biomarkers of diverse cancers in the Human Protein Atlas (https://www.proteinatlas.org) (Uhlén *et al*, 2015), thus confirming their relevance in our study and their potency as biomarker candidates associated to gastric preneoplasia or GC.

#### Short-listing of 15 biomarker candidates from the LC-MS/MS analysis

Based on the analysis of LC-MS/MS data, 15 proteins were short-listed for further confirmation by ELISA, considering both the results of the statistical analyses, their biological functions and their relevance in cancer (Table 2). Statistically, all these proteins were found differentially abundant either in the “GC *vs* preneoplasia” or “GC *vs* heathy” comparison using the “strict” statistical analysis and could thus be biomarker candidates of GC. They were also significantly differentially abundant in at least one other comparison when using the “strict” statistical analysis, except LEP which presents some missing values that exclude it from this analysis. However, LEP was also particularly interesting to test by ELISA, according to the sparse PLS-DA results (Figure 3D). In addition, 7 of these 15 proteins were also differentially abundant between patients affected by preneoplasia and healthy controls in the “relaxed” statistical analysis (Carbonic anhydrase 2 (CA2), DCD, F13A1, Haptoglobin (HPT), Insulin-like growth factor-binding protein complex acid labile subunit (IGFALS), KRT19, and MAN2A1 and could thus be potential biomarkers of gastric preneoplasia.

**Table 2:** List of fifteen proteins from plasma proteomics analysis that were measured by ELISA. Names, KEGG pathways, reactome pathways, and summary informations from the Cancer resource of the Human Protein Atlas (HPA) are reported. The HPA Cancer resource provides information on mRNA expression, including whether a gene is enriched in a particular cancer type (^«^ Cancer enriched ^»^ : at least four-fold higher mRNA level in a particular tissue/cell type compared to any other tissues/cell types ; ^«^ Cancer enhanced ^»^ : at least four-fold higher mRNA level in a particular tissue/cell type compared to the average level in all other tissues/cell types ; or low cancer specificity), and whether it is (^«^ potential ^»^ or ^«^ validated ^»^) prognostic for patient survival in each cancer types from The Cancer Genome Atlas (TCGA).*High expression unfavorable ; **High expression favorab

| Protein name | Uniprot code | Protein description | KEGG pathways | Human Protein Atlas (TCGA) |
| --- | --- | --- | --- | --- |
| **ARG-1** | P05089 | Arginase-1 | hsa01100 Metabolic pathways<br>hsa00220 Arginine biosynthesis | *Cancer enriched*<br>Liver Hepatocellular Carcinoma<br>No prognostic found |
| **ATAD3B** | Q5T9A4 | ATPase family AAA domain containing 3B | hsa03029 Mitochondrial Biogenesis | *Cancer enhanced*<br>Testicular Germ Cell Tumor<br>*Potential prognostic \**<br>Liver Hepatocellular Carcinoma,<br>Kidney Renal Clear Cell Carcinoma. |
| **CA2** | P00918 | Carbonic anhydrase 2 | hsa00910 Nitrogen metabolism<br>hsa01100 Metabolic pathways<br>hsa04971 Gastric acid secretion | *Cancer enriched*<br>Kidney Chromophobe Renal Cell Carcinoma<br>*Potential prognostic\**<br>Lung Squamous Cell Carcinoma |
| **DCD** | P81605 | Dermcidin | hsa09193 Unclassified: signaling and cellular processes | *Cancer enriched*<br>Breast Invasive Carcinoma<br>No prognostic found |
| **F13A1** | P00488 | Coagulation Factor XIII A Chain | hsa09151 Immune system<br>hsa04610 Complement and coagulation cascades | *Low cancer specificity*<br>*Potential prognostic\**<br>Lung Squamous Cell Carcinoma |
| **HPT** | P00738 | Haptoglobin | hsa09181 Protein families metabolism<br>hsa01002 Peptidases and inhibitors Serine peptidases<br>hsa04147 Exosomal proteins of other cancer cells | *Cancer enriched*<br>Liver Hepatocellular Carcinoma<br>*Validated prognostic\**<br>Kidney Renal Clear Cell Carcinoma,<br>Stomach Adenocarcinoma,<br>Lung Adenocarcinoma<br>*Potential prognostic\*\**<br>Hepatocellular Carcinoma |
| **IGFALS** | P35858 | Insulin-like growth factor-binding protein complex acid labile subunit | hsa04935 Growth hormone synthesis, secretion and action<br>hsa04147 Exosomal proteins of haemopoietic cells | *Cancer enhanced*<br>Hepatocellular Carcinoma<br>*Potential prognostic\*\**<br>Hepatocellular Carcinoma,<br>Lung Adenocarcinoma |
| **JUP** | P14923 | Junction plakoglobin | hsa04820 Cytoskeleton in muscle cells<br>hsa05200 Pathways in cancer<br>hsa05202 Transcriptional misregulation in cancer<br>hsa05221 Acute myeloid leukemia<br>hsa05226 Gastric cancer hsa05412 Arrhythmogenic right ventricular cardiomyopathy | *Low cancer specificity*<br>*Potential prognostic\**<br>Hepatocellular Carcinoma<br>*Potential prognostic\*\**<br>Kidney Renal Clear Cell Carcinoma. |
| **KIF20B** | Q96Q89 | Kinesin-like protein KIF20B | hsa04814 Motor proteins<br>hsa04131 Membrane trafficking<br>hsa04812 Cytoskeleton proteins | *Low cancer specificity*<br>*Potential prognostic\**<br>Lung Adenocarcinoma,<br>*Potential prognostic\*\**<br>Kidney Renal Clear Cell Carcinoma<br>*Validated prognostic\**<br>Pancreatic Adenocarcinoma |
| **KRT14** | P02533 | Keratin, type I cytoskeletal 14 | hsa04915 Estrogen signaling pathway<br>hsa05150 *Staphylococcus aureus* infection<br>hsa04147 Exosomal proteins of epithelial, colorectal cancer, melanoma cells | *Cancer enriched*<br>Head and Neck Squamous Cell Carcinoma<br>*Potential prognostic\*\**<br>Breast Invasive Carcinoma<br>*Potential prognostic\**<br>Lung Adenocarcinoma |
| **KRT19** | P08727 | Keratin, type I cytoskeletal 19 | hsa04915 Estrogen signaling pathway<br>hsa05150 *Staphylococcus aureus* infection<br>hsa04147 Exosomal proteins of epithelial, colorectal cancer, melanoma cells | *Low cancer specificity*<br>*Potential prognostic\**<br>Pancreatic Adenocarcinoma<br>*Validated prognostic\**<br>Kidney Renal Clear Cell Carcinoma |
| **LBP** | P18428 | Lipopolysaccharide-binding protein | hsa04064 NF-κB signaling pathway<br>hsa04620 Toll-like receptor signaling pathway<br>hsa04936 Alcoholic liver disease<br>hsa05152 Tuberculosis<br>hsa05417 Lipid and atherosclerosis<br>hsa04147 Exosomal proteins of other cancer cells | *Cancer enriched*<br>Liver Hepatocellular Carcinoma<br>*Validated prognostic\**<br>Kidney Renal Clear Cell Carcinoma<br>*Potential prognostic\**<br>Kidney Renal Papillary Cell Carcinoma |
| **LEP** | P41159 | Leptin | hsa04060 Cytokine-cytokine receptor interaction<br>hsa04080 Neuroactive ligand-receptor interaction<br>hsa04081 Hormone signaling<br>hsa04152 AMPK signaling pathway<br>hsa04630 JAK-STAT signaling pathway<br>hsa04920 Adipocytokine signaling pathway<br>hsa04932 Non-alcoholic fatty liver disease | *Cancer enhanced*<br>Breast Invasive Carcinoma<br>No prognostic found |
| **MAN2A1** | Q16706 | Alpha-mannosidase 2 | hsa00510 N-Glycan biosynthesis<br>hsa00513 Various types of N-glycan biosynthesis<br>hsa01100 Metabolic pathways | *Low cancer specificity*<br>*Potential prognostic\*\**<br>Kidney Renal Clear Cell Carcinoma<br>*Potential prognostic\**<br>Thyroid Carcinoma |
| **S100A12** | P80511 | S100 calcium-binding protein A12 | hsa04990 EF-hand domain-containing proteins > S100 proteins | *Cancer enriched*<br>Head and Neck Squamous Cell Carcinoma<br>*Potential prognostic\**<br>Stomach Adenocarcinoma |

#### ELISA-based validation of 15 biomarker candidates highlighted by LC-MS/MS

ELISAs were used to measure the plasma concentrations of selected biomarker candidates in all samples of the cohort (n=138) (Figure 1). ELISA results confirmed LEP as a potent biomarker, in agreement with the findings from LC-MS/MS analyses. LEP plasma level is differentially abundant between patients with gastritis or preneoplasia *vs* healthy individuals, and patients with preneoplasia *vs* those with gastritis or GC (Figure 4). It is higher in the group of patients with preneoplasia than in the other categories and significantly lower in men than women for all groups, except GC (Figure S3). These findings suggest that plasma LEP concentration could serve as a robust biomarker for identifying preneoplastic lesions, in agreement with previous studies (Capelle *et al*, 2009) (Tas *et al*, 2018).

**Figure 4.**
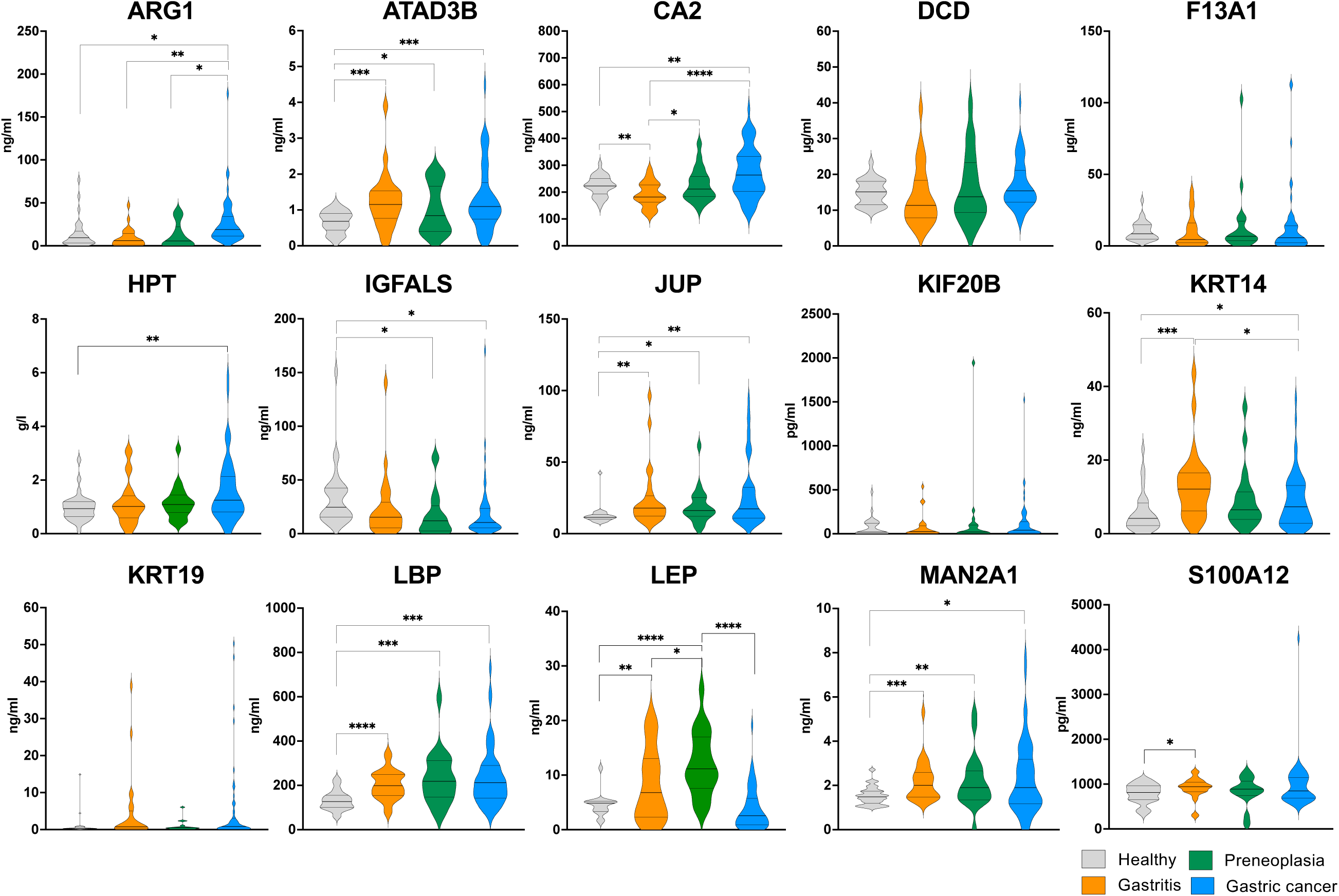
ELISA quantified plasma levels of 15 protein candidates from MS analysis to distinguish patient groups. Protein levels were measured by ELISA commercial assays as indicated in the methods section, on all samples of the cohort. Statistical analysis was performed using Mann-Whithey test. Only significant differential levels are indicated (\*\*\*\**P*<0.0001; \*\*\**P*<0.001; \*\**P*<0.01; \**P*<0.05)

Differential plasma levels between patients with preneoplasia and healthy subjects were also observed for ATAD3B, IGFALS, Junction Plakoglobin (JUP), LBP and MAN2A1 (Figure 4), which along with Arginase-1 (ARG1), CA2, HPT and KRT14 are also differentially abundant between GC patients and healthy individuals. Gender affects the plasma concentration of ATAD3B, JUP and MAN2A1 differentially abundant specifically in women between each group of patients and healthy individuals (Figure S3). When comparing GC patients to healthy individuals, differential plasma levels are observed in women for HPT and in men for IGFALS. Age can also have an impact as in the case of ARG1 more abundant in older GC patients than in others (Figure S4). These 9 proteins with diagnostic properties modulated by age or gender could thus be considered as potent biomarkers of gastric preneoplasia or GC.

Although some of these proteins showed a promising trend, defining specific concentration thresholds for reliably diagnosing a given pathology in individual patients remains challenging.

As reported in previous studies, single protein biomarkers often lack sufficient discriminatory power on their own. However, their diagnostic accuracy can be significantly enhanced when combined with other candidate biomarkers (Bazin *et al*, 2024). Next, we investigated the relationships between the biomarkers concentration and evaluated the added value of multi-marker combinations to improve patient classification.

#### Correlations between plasma concentration levels of the biomarkers

Pearson correlation coefficients analysis between the biomarker concentrations in patients showed that many of them appear strongly significantly different from zero with a p-value threshold of 1% (Figure 5A). Many of the biomarkers are positively correlated. Their plasma concentration evolves globally similarly, as KRT19, CA2 and LBP between them, HPT with DCD, DCD with CA2, JUP with LBP, KIF20B and ARG1 with F13A1, MAN2A1 with ATAD3B. Inversely, MAN2A1 is negatively correlated with IGFALS and CA2, as S100A12 and ATAD3B, or LEP with patient gender (i.e. higher levels in women than men). These observations led to four main protein clusters when using hierarchical clustering based on an absolute Pearson correlation distance (Figure 5B). The first cluster contains KRT19, S100A12, LBP, JUP, CA2, DCD and HPT, the second LEP and gender, the third includes age and *H. pylori* infection status of the patient with ATAD3B, MAN2A1, IGFALS and KRT14, while the fourth is composed of F13A1, KIF20B and ARG1. These different behaviors of biomarkers therefore indicate that not all biomarkers provide the same information. The use of their combination can thus be a positive strategy leading to a more accurate classification of patients.

**Figure 5.**
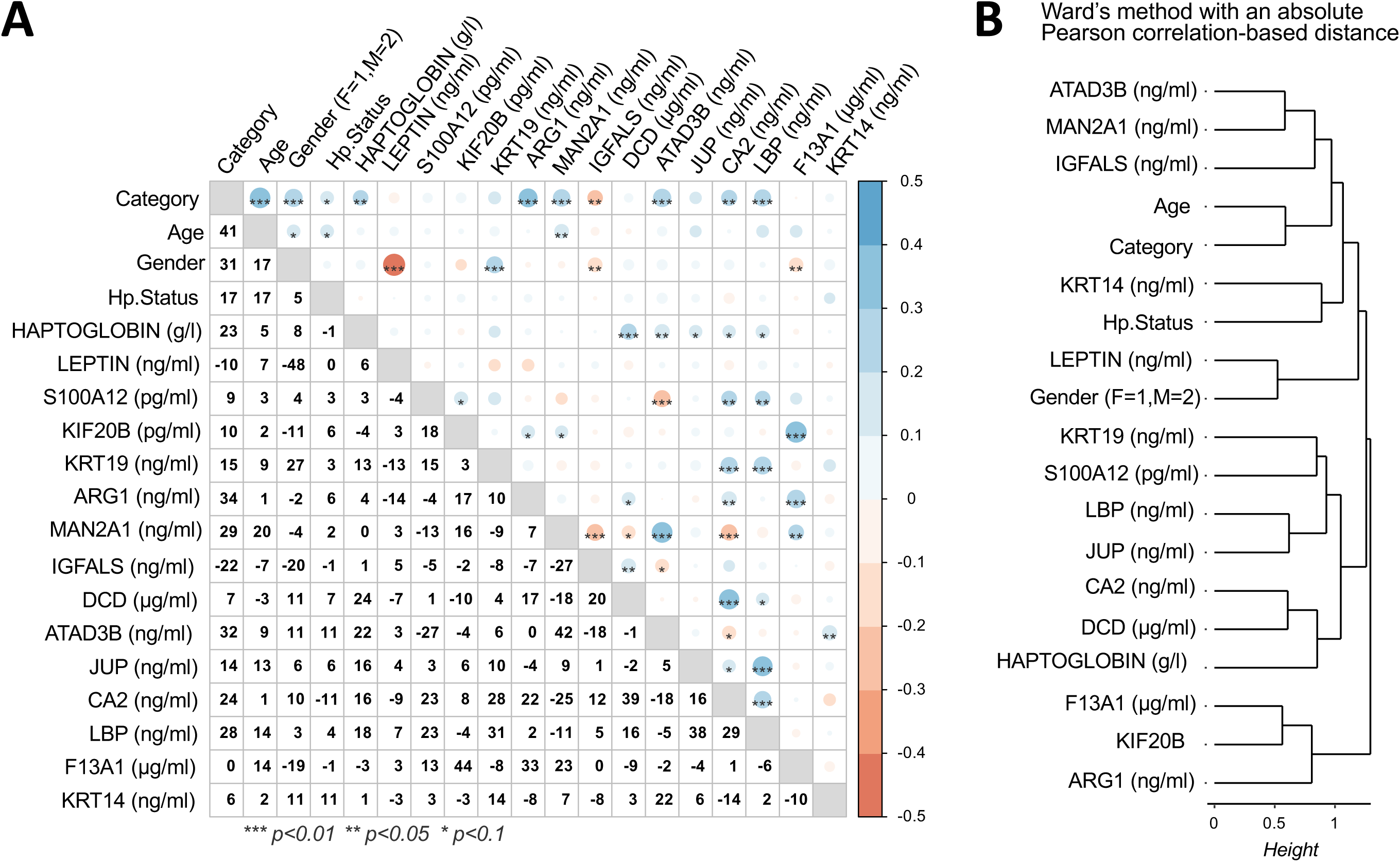
Correlations between biomarker concentrations measured by ELISA across patient samples. A Pearson correlation matrix between plasma concentration levels of the biomarkers. B Hierarchical clustering of biomarkers using a correlation-based distance and the Ward method.

#### Performance of prediction models with single and multiple biomarkers

As described in the Methods section, six types of prediction models were developed using either linear logistic regression or random forests, evaluated through 4-fold cross-validation repeated 10 times (Figure 6A). Models were evaluated based on the mean and standard deviation of the AUROC criterion, as calculated from the test sets of the cross-validation process (Supplementary material additional file 3). We first evaluated each of the 15 ELISA-validated protein biomarkers individually, also including age, gender and *H. pylori* infection status as additional factors that can be used in our prediction models to eventually improve their performance, thus resulting in 18 biomarkers.

**Figure 6.**
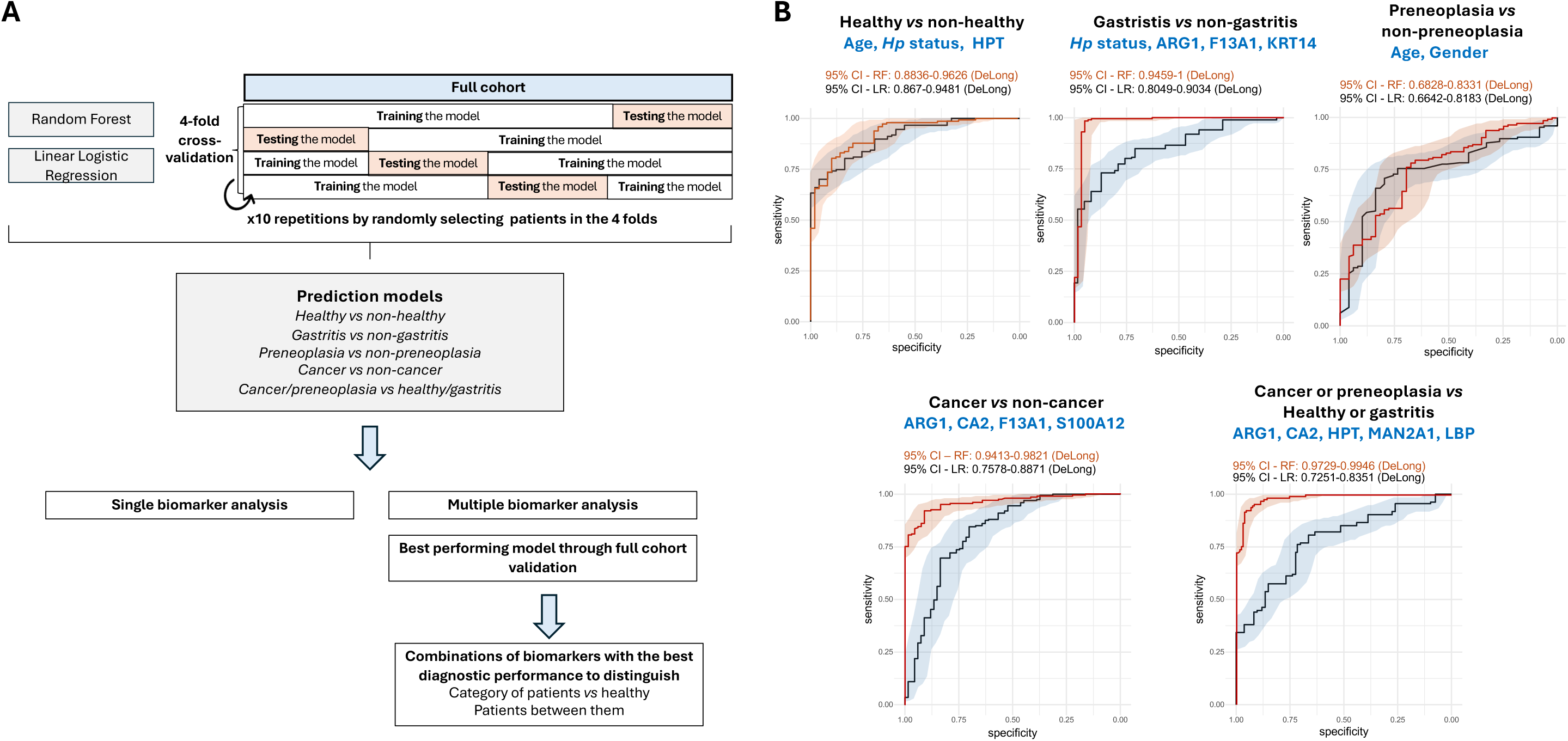
Prediction models with single and multiple biomarkers. A Schematic representation of the 4-fold cross-validation strategy repeated 10 times, used to evaluate prediction models and methods used to identify the combinations of biomarkers with the best mAUROC values. B ROC curves assessed from the full cohort based on models using biomarkers with the best mean AUROC obtained from repeated cross-validations. 95% CI = 95% confidence interval. RF = random forests, LR = linear logistic regression. AUROC values and ROC curves have been obtained using the pROC R package. Confidence intervals have been obtained with the DeLong methods. *Hp: H. pylori*

#### Single biomarker analysis

We first assessed the predictive performance of individual protein biomarkers incorporated into the models. Table 3 reports the mean AUROC (mAUROC) values for all biomarkers across different models, where the reported mAUROC corresponds to the highest value obtained between logistic regression and random forest models (see also Supplementary material additional file 3). For models predicting all patient categories, the best-performing biomarker was the plasma concentration of KRT14 (mAUROC = 63.3%, sd = 4.4%), closely followed by patient age (mAUROC = 63.2%, sd = 4.8%). Notably, age emerged as a crucial factor, yielding the highest mAUROC for several classification tasks, including healthy *vs* non-healthy (mAUROC = 80.1%, sd = 6.9%), preneoplasia *vs* non-preneoplasia (mAUROC = 71.8%, sd = 8.8%), and cancer or preneoplasia *vs* healthy or gastritis (mAUROC = 75.1%, sd = 8.1%). These results reflect the structure of our cohort with healthy individuals younger than patients (Table 1). For cancer vs non-cancer prediction, the best-performing biomarker was the plasma concentration of CA2 (mAUROC = 78.1%, sd = 8.7%). Meanwhile, for gastritis *vs* non- gastritis classification, the highest mAUROC was again obtained with KRT14 (mAUROC = 72.3%, sd = 8.4%). Although these results demonstrate the potent utility of single biomarkers, their moderate mAUROC values indicate that individual biomarkers alone may not be sufficient for robust disease classification.

**Table 3:** Mean and standard deviation of AUROC values for prediction models using single biomarkers. Values represent the highest mean AUROC obtained between logistic regression and random forest models for each prediction task. Green highlighting indicates mAUROC >70% for a single biomarker in a specific classification task.

|  | Healthy vs non-Healthy |  | Gastritis vs non-gastritis |  | Preneoplasia vs non-preneoplasia |  | Cancer vs non-cancer |  | Cancer/Preneoplasia vs Healthy/Gastritis |  | All stages |  |
| --- | --- | --- | --- | --- | --- | --- | --- | --- | --- | --- | --- | --- |
|  | mAUROC | sd | mAUROC | sd | mAUROC | sd | mAUROC | sd | mAUROC | sd | mAUROC | sd |
| Age | 80.1% | 6.9% | 60.2% | 8.1% | 71.8% | 8.8% | 62.2% | 10.0% | 75.1% | 8.1% | 63.2% | 4.8% |
| Gender | 57.0% | 10.9% | 57.7% | 7.7% | 49.6% | 2.4% | 66.0% | 10.4% | 55.4% | 9.5% | 56.6% | 4.0% |
| Hp status | 48.2% | 4.9% | 67.2% | 15.3% | 50.9% | 3.3% | 56.8% | 6.1% | 54.5% | 4.8% | 59.6% | 5.0% |
| DCD | 57.4% | 7.0% | 60.5% | 8.2% | 63.7% | 8.7% | 63.6% | 10.2% | 55.7% | 5.9% | 57.1% | 5.0% |
| IGFALS | 63.4% | 8.7% | 59.6% | 8.0% | 62.5% | 9.3% | 58.3% | 8.0% | 61.2% | 7.8% | 54.1% | 4.9% |
| LEP | 77.0% | 8.5% | 65.1% | 11.4% | 62.4% | 8.9% | 68.4% | 8.8% | 63.2% | 9.1% | 62.7% | 6.5% |
| KRT14 | 63.3% | 7.9% | 72.3% | 8.4% | 61.3% | 8.0% | 58.1% | 6.6% | 55.8% | 5.0% | 63.3% | 4.4% |
| MAN2A1 | 66.0% | 8.0% | 60.3% | 7.8% | 60.7% | 8.3% | 58.8% | 8.2% | 59.5% | 7.4% | 57.8% | 4.8% |
| KIF20B | 57.7% | 8.2% | 57.4% | 7.7% | 60.4% | 7.6% | 62.1% | 8.8% | 57.4% | 7.9% | 49.9% | 5.7% |
| ARG1 | 60.2% | 5.9% | 68.0% | 10.9% | 60.1% | 8.6% | 75.5% | 6.3% | 66.1% | 7.0% | 59.7% | 5.9% |
| F13A1 | 65.6% | 7.8% | 60.5% | 7.1% | 60.0% | 7.4% | 59.1% | 6.6% | 55.3% | 6.0% | 50.4% | 5.1% |
| S100A12 | 57.1% | 8.4% | 59.5% | 8.3% | 59.9% | 8.3% | 61.6% | 10.9% | 58.3% | 6.5% | 53.8% | 5.7% |
| LBP | 70.7% | 6.9% | 60.6% | 9.3% | 59.5% | 7.1% | 68.4% | 9.3% | 67.6% | 6.2% | 61.1% | 4.4% |
| ATAD3B | 61.3% | 8.4% | 61.0% | 7.9% | 59.5% | 8.0% | 59.7% | 8.2% | 57.3% | 8.4% | 50.6% | 7.4% |
| JUP | 71.4% | 8.1% | 58.6% | 7.7% | 59.3% | 6.8% | 63.2% | 9.9% | 65.2% | 6.6% | 58.8% | 5.4% |
| CA2 | 64.9% | 8.6% | 63.9% | 10.8% | 58.8% | 7.8% | 78.1% | 8.7% | 69.4% | 7.7% | 62.0% | 6.1% |
| HPT | 62.0% | 7.9% | 61.1% | 8.9% | 58.5% | 6.2% | 64.1% | 10.0% | 64.8% | 8.1% | 55.6% | 4.9% |
| KRT19 | 63.3% | 9.5% | 61.0% | 9.3% | 57.7% | 8.5% | 60.4% | 7.7% | 56.0% | 7.2% | 53.3% | 3.8% |
| Max mAUROC | 80.1% | 6.9% | 72.3% | 8.4% | 71.8% | 8.8% | 78.1% | 8.7% | 75.1% | 8.1% | 63.3% | 4.4% |
| Min mAUROC | 48.2% | 4.9% | 57.4% | 7.7% | 49.6% | 2.4% | 56.8% | 6.1% | 54.5% | 4.8% | 49.9% | 5.7% |
| Biomarker with best mAUROC | Age |  | KRT14 |  | Age |  | CA2 |  | Age |  | KRT14 |  |

#### Multiple biomarker analysis

To improve predictive accuracy, we examined models incorporating multiple biomarkers using the same methodology (Figure 6A). Given that the number of possible biomarker combinations increases exponentially with the number of biomarkers included (Table 4), we limited our analysis to a maximum of six biomarkers selected from the 18 candidates, resulting in a total of 31,179 unique tested biomarker combinations (Supplementary material additional file 3). As reported in Table 4, models that integrated multiple biomarkers outperformed those using a single one, achieving higher mAUROC values. However, adding more biomarkers did not always lead to better performance, as none of the highest mAUROC values were observed in models containing six biomarkers. For models predicting all patient categories, the best-performing model achieved a mAUROC of 77.9% (sd = 7.7%) and incorporated age, gender, *H. pylori* status and the plasma concentration of ARG1 and KRT14. A different model, designed to predict cancer or preneoplasia *vs* healthy or gastritis, yielded an even higher mAUROC of 83.9% (sd = 5.1%) and was based on the plasma concentrations of ARG1, HPT, CA2, MAN2A1, and LBP. HPT also emerged as a key biomarker to distinguish between healthy and pathological states, as combining with age and *H. pylori* status, this model achieved the highest mAUROC of 88.0% (sd = 5.3%). For cancer *vs* non-cancer prediction, the top-performing model included the plasma concentrations of ARG1, CA2, F13A1 and S100A12, achieving a mAUROC of 85.3% (sd = 6.4%). The best model for gastritis *vs* non-gastritis classification combined *H. pylori* status with the plasma concentrations of ARG1, KRT14, and F13A1, yielding a mAUROC of 82.2% (sd = 9.2%). For preneoplasia *vs* non- preneoplasia prediction, the strongest model was based on the patient age and gender (mAUROC = 75.7%, sd = 8.9%), closely followed by a model that also incorporated the plasma concentration of KRT14 (mAUROC = 74.4%, sd = 10.6%). Thus, predicting preneoplasia *versus* other categories appears more challenging even if the highest mAUROC values exceed 50%, indicating interesting predictive capability.

**Table 4:** Mean AUROC values and standard deviations for prediction models incorporating multiple biomarkers. For each classification task, the highest performing biomarker combination is highlighted in green, along with its corresponding mAUROC value. This table represents only the highest mAUROC obtained between logistic regression and random forest models for each classification task and number of biomarkers in the models.

|  | Number of biomarkers in the prediction model |  |  |  |  |  |  |  |  |  |  |  |
| --- | --- | --- | --- | --- | --- | --- | --- | --- | --- | --- | --- | --- |
|  | 1 |  | 2 |  | 3 |  | 4 |  | 5 |  | 6 |  |
|  | mAUROC | sd | mAUROC | sd | mAUROC | sd | mAUROC | sd | mAUROC | sd | mAUROC | sd |
| Healthy vs non-healthy | 80.1% | 6.9% | 87.7% | 5.0% | 88.0% | 5.3% | 87.4% | 6.9% | 86.2% | 7.9% | 85.2% | 8.2% |
|  | Age |  | Age, *Hp status* |  | Age, *Hp status*, HPT |  | Age, *Hp status*, HPT, KRT19 |  | Age, Gender, *Hp status*, HPT, KRT19 |  | Age, Gender, *Hp status*, ARG1, F13A1 |  |
| Gastritis vs non-gastritis | 72.3% | 8.4% | 81.4% | 7.3% | 80.6% | 7.5% | 82.2% | 9.2% | 79.6% | 9.2% | 78.0% | 8.0% |
|  | KRT14 |  | *Hp status*, KRT14 |  | *Hp status*, F13A1, KRT14 |  | *Hp status*, ARG1, KRT14, F13A1 |  | *Hp status*, ARG1, KRT14, F13A1, KIF20B |  | *Hp status*, HPT, KIF20B, ARG1, F13A1, KRT14 |  |
| Preneoplasia vs non-preneoplasia | 71.8% | 8.8% | 75.7% | 8.9% | 74.4% | 10.6% | 73.4% | 10.8% | 72.7% | 7.9% | 70.5% | 11.3% |
|  | Age |  | Age, Gender |  | Age, Gender, KRT14 |  | Age, Gender, KRT14, ARG1 |  | Age, Gender, KRT14, KRT19, F13A1 |  | HPT, Leptin, S100A12, KIF20B, ATAD3B, CA2 |  |
| Cancer vs non-cancer | 78.1% | 8.7% | 83.0% | 9.4% | 85.2% | 7.0% | 85.3% | 6.4% | 85.0% | 5.7% | 84.9% | 7.4% |
|  | CA2 |  | CA2, Leptin |  | ARG1, CA2, S100A12 |  | ARG1, CA2, F13A1, S100A12 |  | Gender, ARG1, CA2, MAN2A1, IGFALS |  | Age, Gender, ARG1, CA2, MAN2A1, IGFALS |  |
| Cancer/preneoplasia vs healthy/gastritis | 75.1% | 8.1% | 76.7% | 7.7% | 78.8% | 6.8% | 81.2% | 6.8% | 83.9% | 5.1% | 81.4% | 6.3% |
|  | Age |  | Age, Leptin |  | Age, HPT, ARG1 |  | ARG1, CA2, HPT, LBP |  | ARG1, CA2, HPT, MAN2A1, LBP |  | ARG1, CA2, HPT, MAN2A1, LBP, DCD |  |
| All stages | 63.3% | 4.4% | 71.6% | 5.4% | 74.8% | 4.2% | 76.2% | 6.9% | 77.9% | 7.7% | 75.6% | 6.3% |
|  | KRT14 |  | Age, *Hp status* |  | Age, Gender, *Hp status* |  | Age, *Hp status*, ARG1, KRT14 |  | Age, Gender, *Hp Status*, ARG1, KRT14 |  | Age, Gender, *Hp Status*, ARG1, KRT14, F13A1 |  |
| Number of tested combinations | 18 |  | 153 |  | 816 |  | 3060 |  | 8568 |  | 18 564 |  |

#### Full cohort validation

Finally, we assessed the best performing models on the full cohort to obtain final performance estimates. Using the combinations of biomarkers displaying the best mAUROC values with repeated cross-validations, we assessed their performance to classify patients on the full cohort using linear logistic regression and random forests (Figure 6A). Notably, the AUROC values for the models classifying cancer *vs* non-cancer, and cancer/preneoplasia *vs* healthy/gastritis, were estimated superior to 94.1% and 97.3%, respectively, using random forests (lower bounds of 95% confidence intervals) (Figure 6B). This demonstrates that the combinations of short-listed biomarkers corresponding to ARG1, CA2, F13A1 and S100A12 for the distinction cancer *vs* non cancer and ARG1, CA2, HPT, MAN2A1, and LBP to distinguish cancer or preneoplasia *vs* healthy or gastritis, enable near-perfect classification of patients for these tasks on the full cohort.

## Discussion

GC is a multifactorial disease often diagnosed at an advanced stage, limiting the efficacy of therapeutic interventions and contributing to its poor prognosis. Currently, diagnosis relies on upper endoscopy, an invasive and resource-intensive procedure that requires specialized equipment and highly skilled clinicians to accurately identify gastric preneoplasia, particularly DYS which need a periodical surveillance (Kim *et al*, 2020)(Young *et al*, 2021). A critical gap is the lack of non-invasive diagnostic methods for early detection of GC. In this context, blood-based biomarkers represent a promising alternative to overcome the limitations of endoscopic screening and to reduce the global burden of GC.

In this study, we used bottom-up LC-MS/MS-based proteomics of plasma samples from patients with gastric lesions at various stages of the carcinogenesis cascade, followed by ELISA validation, to identify and confirm plasma protein biomarkers associated with gastric preneoplastic and cancerous lesions. Our results provide compelling evidence that specific protein concentration profiles in plasma can serve as reliable indicators of disease stage and may be combined to develop non-invasive diagnostic models capable of identifying individuals at high risk of GC, even at asymptomatic stages.

Our LC-MS/MS analyses highlighted numerous candidate biomarkers across patient categories (408 differentially abundant proteins in the “relaxed” statistical analysis and 236 in the “strict” analysis). These proteins are involved in pathways known to be dysregulated in cancer, as annotated in KEGG (https://www.genome.jp/kegg/pathway.html) and the Human Protein Atlas (https://www.proteinatlas.org) databases, including proteins related to signaling and cellular processes (DCD, IL1RAP, KRT14, KRT19, S100A12, SAA1/SAA2, SVEP1, TFRC), innate immunity and immune response (F13A1, IGHG1, LBP), cell motility and membrane trafficking (CAPZA1, KIF20B), glycan biosynthesis (FUCA2, MAN2A1), digestive and cancer-related systems (CA2, JUP, LEP, PRCP, PRSS3), metabolism, cell energy and growth (ARG1, ATAD3B, ERAP2, GLPD1, HPT, IGFALS). Most of these proteins are recognized as potential prognostic biomarkers in liver (TFRC and GPLD1), renal (IGHG1, PRCP, SAA1/2 and IL1RAP) and digestive (PRSS3) cancers in the Human Protein Atlas database. Importantly, the GC proteomics database (GCPDB) (Ramesh *et al*, 2024), has previously reported differential levels of IGFALS, LBP, S100A12, F13A1, SAA1/SAA2, PRCP, and IGHG1 in plasma/serum proteome MS analysis of GC patients at various stages, further validating the relevance of our findings.

From our discovery set, fifteen proteins were short-listed, among which several emerged as promising biomarkers after validation by ELISA in the entire cohort of 138 subjects (including 106 patients). LEP was particularly noteworthy, displaying significant differential abundance across several patient groups, with highest levels in patients with preneoplasia. Given its diverse functions in the digestive system, including the regulation of immune response, cell proliferation and tissue repair, lipid and glucose metabolism (Kim *et al*, 2021), LEP represents a biomarker of special interest for gastric preneoplastic and carcinogenic processes. Previous studies have already associated higher LEP levels with gastric inflammation and IM (Capelle *et al*, 2009), supporting our findings. Additional proteins, ATAD3B, IGFALS, JUP, LBP, and MAN2A1 also displayed significantly higher plasma concentration in the preneoplasia group compared to healthy individuals, suggesting them as early markers of gastric mucosal transformation.

When comparing GC patients to healthy individuals, nine proteins (ARG1, ATAD3B, CA2, HPT, IGFALS, JUP, KRT14, LBP and MAN2A1) displayed significantly altered plasma levels. Many of these identified biomarkers have established roles in carcinogenesis. ARG1 modulates oncogenic signaling pathways such as PI3K/AKT/mTOR, STAT3 and MAPK (Niu *et al*, 2022). It has also been mentioned as enriched in hepatocellular carcinoma (Wang *et al*, 2020). ATAD3B, a mitochondrial membrane-bound protein, has been associated with tumorigenesis (Li *et al*, 2012) and suggested as potential prognostic biomarker for hepatocellular carcinoma patients (Liu *et al*, 2019). CA2, a metalloenzyme involved in gastric acid secretion (Kivelä *et al*, 2005), helps regulate pH, which is vital for cancer cell survival in acidic tumor microenvironments. Its low expression correlates with poor prognosis in GC patients (Hu *et al*, 2014). HPT, a hemoglobin-binding glycoprotein associated with inflammatory response, shows aberrant glycosylation in GC, suggesting it as a signature molecule for GC diagnosis and detection (Jeong *et al*, 2020). It has also been previously proposed as biomarker to predict colorectal cancer and tumor progression (Ghuman *et al*, 2017). IGFALS, crucial for the stability of circulating insulin growth factors, is down-regulated in hepatocellular carcinoma and associated with poor prognosis (Xu *et al*, 2024). JUP, a cell-cell junction protein and transcription factor, exhibits loss correlated with GC malignancy and poor prognostics (Chen *et al*, 2021). KRT14, a member of the keratin family of intermediate filament proteins, could be an indicator of the remodeling of the gastric epithelium, promoting tumor growth and invasion (Zheng *et al*, 2016). LBP has been linked to GC pathogenesis and proposed as both a prognostic biomarker (Lv *et al*, 2025) and a diagnostic biomarker for liver metastasis in GC (Xie *et al*, 2023).

MAN2A1, involved in N-linked oligosaccharide processing in the Golgi apparatus, may contribute to the metabolic reprogramming and altered glycosylation characteristic of cancer cells. It has been proposed as a potential biomarker significantly related to prognosis and lymph node metastasis in colorectal cancer patients (Wang *et al*, 2022).

While individual biomarkers showed encouraging trends, the heterogeneity in expression patterns and inter-patient variability prevented accurate classification of patient categories based on single protein measurements. Our correlation analysis revealed distinct biomarker clusters, suggesting that combining multiple markers could enhance classification accuracy and disease stage differentiation. From the prediction models-based analysis two panels proved particularly effective: ARG1, CA2, HPT, MAN2A1 and LBP (mAUROC: 83.9%) for differentiating patients with cancer or precancerous lesions from healthy individuals or patients with gastritis; and ARG1, CA2, F13A1 and S100A12 (mAUROC: 85.3%) for differentiating GC from non-cancer patients. ARG1 and CA2 appeared in both panels, reinforcing their significance as GC biomarkers, supported by their high individual mAUROC values (75.5% and 78.1%, respectively). Interestingly, while S100A12 and F13A1 did not show significant individual differences between GC patients and healthy individuals, they enhanced diagnostic performance when combined with ARG1 and CA2. S100A12, a member of calgranulin protein family, is related to innate immunity and has been proposed as a prognostic factor in GC (Li *et al*, 2016), while F13A1 has been associated with tumor metastasis in lung cancer and poor prognosis in (Ercan *et al*, 2021) several cancers (Lee *et al*, 2013) (Raval *et al*, 2018) (Liu *et al*, 2025). From a biological point of view, the combination of these different proteins is relevant, reflecting early step of gastric carcinogenesis and tumor development-related mechanisms. The inclusion of proteins like ARG1 and CA2 in our most effective predictive models suggests that metabolic reprogramming plays a crucial role in GC development. ARG1, by depleting arginine, may create an immunosuppressive microenvironment facilitating tumor immune evasion, while alterations in CA2 could reflect disruption in pH regulation critical for cancer cell adaptation. The inflammatory proteins in our panels, including S100A12 and LBP, may form part of interconnected signaling networks linking inflammation, immune modulation, and metabolic adaptation driving the progression from gastritis to preneoplasia and cancer. The presence of HPT for distinguishing cancer/preneoplasia from healthy/gastritis states could further highlight the significance of disrupted iron metabolism and acute phase response pathways in gastric carcinogenesis.

While our identified biomarkers show promise for GC detection, it is important to acknowledge that many are not exclusive to gastric pathology. Several proteins in our panels, have been associated with other cancer types or inflammatory conditions as ARG1 (Wang *et al*, 2020) , HPT (Ghuman *et al*, 2017), IGFALS (Xu *et al*, 2024) and MAN2A1 (Wang *et al*, 2022). However, the strength of our approach lies in the specific combination of these biomarkers rather than individual proteins. Therefore, the multi-protein signatures we identified likely reflect the unique molecular patterns of gastric carcinogenesis and precancerous progression. This suggests that while individual proteins may lack specificity, their coordinated alterations create distinctive signatures that may be more specific to gastric pathology. Future studies comparing these signatures across different gastrointestinal and inflammatory conditions would further clarify their diagnostic specificity.

Among our classification tasks, predicting preneoplasia versus other categories proved particularly challenging, with relatively lower mAUROC values than for the other tasks. Although the best models achieved mAUROC values above 74% for this task, indicating promising predictive capability, their performances suggest additional biomarkers or alternative modeling approaches may be necessary for this category. The lower diagnostic performance observed in this task could be attributed to the biological heterogeneity within the preneoplasia group, which includes patients with AG, IM and DYS.

Recent studies have explored various protein combinations for GC detection, such as the panel reported by Shen et al. (2019), but few have specifically addressed the continuum from gastritis to preneoplasia to cancer as comprehensively as our approach. Our multi-protein panels achieved significantly higher diagnostic performance, than existing serological cancer biomarkers (Feng *et al*, 2017) with mAUROC values exceeding 85% for cancer detection and 83% for identifying patients with cancer or preneoplasia using cross-validations. However, while our study reported promising results, with biomarker panels achieving AUROC values exceeding 94% for distinguishing GC from non-GC on the full cohort, some limitations should be acknowledged. First, the age disparity between our healthy control group (mean 42 years) and disease groups (59-70 years) represents an inherent challenge in biomarker studies for age-associated diseases like GC. To address this potential confounder, we explicitly incorporated age into our predictive models, which allowed to distinguish age-related effects from disease-specific protein alterations. Notably, while age emerged as a strong predictor in several classification tasks (mAUROC 80.1% for healthy vs non-healthy), our multi-protein panels maintained high predictive performance even when controlling for age effects. For instance, the combination of ARG1, CA2, F13A1, and S100A12 achieved high discrimination of cancer vs non-cancer cases (mAUROC : 85.3%) that cannot be attributed to age differences alone, as evidenced by the lower performance of age as a single predictor (mAUROC : 62.2%) for this specific classification. This suggests that our identified protein signatures reflect disease-specific processes rather than merely age-associated changes in plasma proteome composition. Second, while our cohort (138 participants) provided sufficient power to identify several promising biomarkers, larger studies are necessary to strengthen these findings, including external validation in independent cohorts with diverse demographic and clinical characteristics.

From a public health perspective, our findings have far-reaching implications. Implementation of blood-based testing could substantially improve screening participation in both high-risk populations and resource-limited settings where endoscopic capacity is insufficient. In countries with established screening programs (i.e. Japan, South Korea), our approach could help refine risk stratification, potentially reducing unnecessary endoscopies while maintaining high detection rates. Importantly, in Western countries with no screening strategies for GC, this test could enable more targeted surveillance particularly for individuals not identified as high-risk, currently detected at late stages. This is even more important, considering the concerning and recent rise in GC cases among younger populations.

## Conclusion

Our study presents a comprehensive approach for the early detection of GC and its precursor lesions by leveraging plasma proteomics. Based on predictive models achieving high classification performance using a minimal set of proteins, combinations of biomarkers with high diagnosis accuracy have been identified for distinguishing cancer from non-cancer (AUROC>94%) and cancer or preneoplasia from healthy or non-atrophic gastritis cases (AUROC>97%). These findings pave the way to future development of a non-invasive and highly predictive blood test as an initial screening tool to identify at the earliest individuals at high-risk of GC. This diagnostic test could also be useful for the follow-up of patients previously detected with preneoplasia as well as to follow the recurrence/ remission of patients under anticancer treatment, thus optimizing patient care and survival.

### Methods and Protocols

#### Study population

The study cohort consists of 138 participants. It includes 32 healthy asymptomatic volunteers recruited at the Institut Pasteur clinical investigation and biomedical research support unit (ICAReB-Clin). They were confirmed for their *H. pylori*-negative serology using a commercial ELISA (Serion/diagnostic, Denmark). The groups of patients included 26 cases of non-AG referred as “gastritis”, 20 of AG/preneoplasia, referred as the “preneoplasia” group, including patients with AG either with IM or DYS or not (Table 1). Patients with gastritis and preneoplasia were recruited at the unit of Hepato-Gastroenterology, AP-HP, A. Paré Hospital, Boulogne-Billancourt, France. The 60 patients with GC (stages III and IV) were enrolled from the unit of Digestive Oncology, AP-HP, European Hospital G. Pompidou (HEGP), Paris, France. All patients were previously diagnosed through upper endoscopy combined with gastric biopsies and anatomopathology analysis. All patients were adults, they had not received antibiotherapy and had not been treated with bismuth compounds, proton pump inhibitors or non-steroidal anti-inflammatory drugs for at least two weeks prior to the study. Each patient was informed about the study and provided written consent. For each patient, 10 ml of blood were collected and transferred to Leucosep^®^ tubes (Dutscher, France) for plasma isolation according to recommendation of the supplier.

The study was approved by the Institut Pasteur Clinical Research Coordination Office (Ref 2013- 29), the French advisory committee on information processing in Health Research (Ref 14.202) and the French data protection authority: Commission Nationale de l’Informatique et des Libertés (CNIL) (Ref: 2017-204).

### Identification and quantification of plasma proteomes using LC-MS/MS

#### Plasma samples for LC-MS/MS analysis

As indicated in Table 1, plasma proteome identification and quantification were performed on samples from a subset of cohort patients with gastritis (n=10), preneoplasia (n=9), GC (n=10), and healthy controls (n=10). The most abundant proteins were first depleted from plasma, and the remaining protein samples underwent proteolytic digestion, as detailed in the Supplementary material. Next, proteins identification and quantification were performed using Data-Independent Acquisition (DIA), based on a spectral library obtained through Data-Dependent Acquisition (DDA).

#### LC-MS/MS analysis

Mass spectrometry-based plasma proteomics were conducted in two steps. The 1^st^ step was dedicated to the spectral library generation from DDA using the fractionated “pool” sample. The 2^nd^ step was defined to specifically identify and quantify plasma proteome for each patient group from DIA. For this purpose, a nanochromatographic system (Proxeon EASY-nLC 1200 - Thermo Fisher Scientific, Waltham, Massachusetts, USA) was coupled online to a Q Exactive^TM^ HF Mass Spectrometer (Thermo Fisher Scientific) using an integrated column oven (PRSO-V1 - Sonation GmbH, Biberach, Germany). For each sample, 1 μg of peptides was injected onto a home-made C18 column. Peptides elution and mass spectra acquisition were performed as described in the Supplementary material. Data processing for proteins identification and quantification including building of the spectral library and data analysis for DIA method are reported in the Supplementary material.

### Statistical analysis of large-scale LC-MS/MS proteomic data for biomarker identification

#### Multivariate analysis of the MS-based proteomics data

Pairwise correlation analysis, hierarchical clustering and PLS-DA have been performed to highlight global similarities between quantified proteomes of plasma samples. The correlation matrix represents the Pearson correlation coefficients between each pair of samples computed using all complete pairs of intensity values measured in these samples. The hierarchical cluster analysis has been conducted via multiscale boostrap resampling (1000 bootstrap replications) with the Ward’s method and a correlation-based distance measure thanks to the *pvclust* function of the *pvclust* R package (Suzuki & Shimodaira, 2006), after log2 transformation of the intensities, imputation of the missing values with the *impute.slsa* function of the *imp4p* R package (Gianetto *et al*, 2020) and a normalization using a sample-median centering method inside conditions. PLS-DA using 5 components was performed using the *mixOmics* R package (Rohart *et al*, 2017) to assess whether the quantified proteomes differ among the four patient groups (Healthy, Gastritis, Preneoplasia, and GC).

#### Identification of potential biomarkers from MS-based proteomics data

To identify potential protein biomarkers, pairwise differential analyses (e.g. comparisons of one category of patient *vs* another) were conducted, as well as a multivariate sparse PLS-DA.

To identify proteins that were more abundant in one condition than another, quantified intensities obtained using Spectronaut were compared. Proteins were retained for further statistical analysis only if they had at least 2 quantified intensity values (for the "relaxed" analysis) or 9 quantified intensity values (for the "strict" analysis) in at least one of the two conditions being compared. Proteins that were quantified in one condition but not in the other were set aside, as they could be directly considered as potential biomarkers. Following this initial filtering, the intensities of the remaining proteins were log-transformed (log2). Next, intensity values were normalized by median centering within a same category of patients (section 3.5 in (Giai Gianetto, 2023)). Missing values were imputed using the *impute.slsa* function from the *imp4p* R package (Gianetto *et al*, 2020). Proteins with a fold-change (FC) below 2 (i.e., absolute log₂(FC) < 1) were considered not significantly differentially abundant. Statistical testing was then performed using a *limma* t-test, implemented in the *limma* R package (Ritchie *et al*, 2015). To control for multiple testing, an adaptive Benjamini-Hochberg procedure was applied to the resulting p-values using the *adjust.p* function from the *cp4p* R package (Giai Gianetto *et al*, 2016). The robust method described in (Pounds & Cheng, 2006) was used to estimate the proportion of true null hypotheses among the statistical tests. Proteins with an adjusted p-value below a false discovery rate (FDR) of 1% were considered significantly differentially abundant. Ultimately, proteins identified as potential biomarkers were those that emerged from this multiple testing analysis, along with those quantified in one condition but not in the other. The lists of potential biomarkers obtained from the different comparisons were next compared using Venn diagrams to highlight proteins that are differentially abundant in several comparisons.

A sparse PLS-DA was performed alongside pairwise comparisons of patient categories. Unlike the previous approach, which compares patient groups two by two, sparse PLS-DA aims to distinguish all groups simultaneously using a restricted set of proteins. For this analysis, only proteins with at least two quantified values in all patient categories were considered. Similar to the differential analyses, the intensities of the selected proteins were log-transformed (log2), normalized by median centering within a same category of patients (section 3.5 in (Giai Gianetto, 2023)) , and missing values were imputed using the *impute.slsa* function from the *imp4p* R package (Gianetto *et al*, 2020). Finally, sparse PLS-DA was applied, with a constraint of selecting only 10 proteins per component, using the *mixOmics* R package (Rohart *et al*, 2017). We employed multiple complementary analytical strategies to ensure robust biomarker identification. The « relaxed » analysis (requiring at least two quantified values in patient categories) maximized sensitivity for discovery of potential biomarkers, while the « strict » analysis (requiring at least nine quantified values) prioritized reproducibility and reliability. This dual approach allowed us to balance biomarker discovery with confidence in the findings. The sparse PLS-DA analysis provided a complementary multivariate perspective, identifying proteins capable of distinguishing all patient groups simultaneously rather than through pairwise comparisons. This multi-faceted statistical strategy strengthens confidence in our findings and provides more reliable performance estimates than single validations would allow.

### Validation of LC-MS/MS-identified proteins using Enzyme Linked Immunosorbent Assays (ELISA), statistical analysis and prediction models

To validate the protein biomarker candidates deduced from the analysis of the LC-MS/MS data, the plasma concentration of 15 short-listed proteins was measured across the full cohort of patient plasma samples (138 samples) using ELISA assays according to the manufacturer’s instructions (MyBiosource, USA and R&D systems, USA) (Supplementary material- Table S1).

The final selection of 15 proteins for ELISA validation was based on (1) statistical significance in the LC-MS/MS analyses, particularly proteins showing differential abundance in multiple comparisons; (2) biological relevance to cancer processes as documented in literature and pathway databases. Statistical analysis of the concentration levels was performed using the Student’s t-test or Mann-Whitney test. Results were considered significant if p<0.05.

#### Correlation analysis of measured concentrations

The measured concentrations were analyzed using Pearson correlation coefficients between biomarkers, using all complete pairs of concentration values measured in these samples. A hierarchical cluster analysis has been conducted with the Ward’s method and a correlation-based distance measure derived from the correlation coefficients. This approach was used to identify biomarkers displaying similar concentration patterns across patients.

#### Estimation of prediction models to distinguish patient categories from measured concentrations

To estimate prediction models for the different pathologies, we used a 4-fold cross validation strategy repeated 10 times. It consists of randomly splitting the set of patients into four sets while preserving the proportion of patients associated to the different pathologies in each as in the original dataset. Three sets of patients over four are then used alternatively to create a training dataset used to estimate the prediction model, while the fourth set is used to assess the performance of the estimated model (test dataset). An up-sampling technique was used to duplicate patients of the minority categories with replacement to tackle the imbalance of the categories in the training dataset in view to improving model performance. Finally, each model provides an estimated probability that a patient is affected by a pathology from a set of biomarkers. We assessed the results of the estimated prediction models on the remaining set of patients (test dataset) using AUROC. For each model, we randomly repeated 10 times the 4-fold cross validation strategy to assess the variations of results. AUROC values and ROC curves have been obtained using the pROC R package (Robin *et al*, 2011). Confidence intervals have been obtained with the DeLong methods (DeLong *et al*, 1988).

Different types of prediction models were estimated. Firstly, we estimated a model predicting all categories of patients (i.e. predicting whether the patient is suffering from GC, preneoplasia, gastritis or if he/she is healthy). Second, we estimated four binary models specific to each category (i.e. predicting healthy *vs* non-healthy, cancer *vs* non-cancer, etc.). Third, we evaluated a model grouping GC and preneoplasia in one category and healthy and gastritis in another (i.e. predicting GC/preneoplasia *vs* healthy/gastritis). This approach results in six types of prediction models, each of them is estimated by both linear logistic regression and using random forests, leading to twelve estimated models for each combination of biomarkers.

## Supporting information

Supplementary material (Add. file 1, file 2, file 3, Supplementary file)

## Acknowledgements

We thank Pascale Bellamy from the clinical research coordination pole at the Institut Pasteur Paris for her implication in regulatory requirements and cohort setting up. We also would like to thank Sébastien Quesney from the Institut Pasteur Research Application and Industrial Relations Department for his interest in the study and helpful discussions. We are also grateful to the medical staff from the Unit of Hepato-Gastroenterology at the A. Paré hospitals, Boulogne Billancourt and the Unit of Gastroenterology and Digestive Oncology at the Georges Pompidou European hospitals, Paris for their implication in the study and samples handling. We also thank all the patients that accepted to participate to this study.

This work was supported by grants from ACIP (Pasteur International Concerted Action) grant ACIP2015-10) from Institut Pasteur Paris, the Odyssey Reinsurance Company and Program Flash Maturation (Project INNOV-80-20) from the Institut Pasteur Research Application and Industrial Relations Department to ET.

## Author contributions

The authors listed above all contributed to this manuscript and approved the final submitted version.

QGG conceptualized, designed and performed the analysis; wrote the draft. VM designed and performed ELISA experiments and analysis. TD conceptualized, designed and performed the proteomic analyses. KN contributed to analyses. AZ, OC, IT, SP, MNU, CJ, JT, DL recruited patients, contributed to plasma collection and cohort characterization. NJ contributed to regulatory procedures and cohort set-up. DL conceptualized and designed the initial project. MM conceptualized and designed the proteomic analyses. ET conceptualized and designed the initial project, contributed to the analyses and wrote the draft. TD, KN, MM, DL contributed to critical revision of the manuscript. MM, ET supervised the study.

## Data availability

The mass spectrometry proteomics data have been deposited to the ProteomeXchange Consortium via the PRIDE partner repository with the dataset identifier PXD062318.

https://www.ebi.ac.uk/pride

## The Paper Explained

### PROBLEM

Gastric cancer (GC) still remains of public health concern, mainly associated with a poor prognosis due to late detection, with 5-year survival rates of only 15-20% in Western countries compared to 60% in some Asiatic countries where screening programs have been implemented. Current gold standard diagnosis relies on upper endoscopy, which is invasive, costly, and has limited sensitivity for detecting gastric preneoplasia such as dysplasia. There is an urgent need for non-invasive, blood-based biomarkers capable of detecting both GC and its precursor lesions, which could enable earlier intervention and significantly improve patient outcomes and reduce the global burden of this cancer..

## RESULTS

Using a two-phase approach of mass spectrometry discovery followed by ELISA validation in 138 participants, we identified effective plasma protein biomarker panels for gastric preneoplasia and GC detection. Two multi-protein combinations showed exceptional performance: (1) ARG1, CA2, F13A1, and S100A12 distinguished cancer from non-cancer cases with AUROC >94%, and (2) ARG1, CA2, HPT, MAN2A1, and LBP differentiated cancer/preneoplasia from healthy/gastritis cases with AUROC >97% in our full cohort. The proteins represent diverse cancer-related biological processes, including metabolism (ARG1, CA2), inflammation (S100A12, LBP), and iron homeostasis (HPT), providing a comprehensive signature of gastric pathology.

## IMPACT

This blood-based biomarkers discovery approach could lead to important improvement for GC screening and its early detection. By providing a simple blood test with high sensitivity and specificity, screening coverage could significantly expand, particularly in regions with limited endoscopic resources. Implementation could occur as (i) an initial risk assessment tool to identify candidates for endoscopy , (ii) as a complementary test to improve risk stratification, and (iii) to monitor patients following a cancer treatment protocol to survey the recurrence/remission thus optimizing patient management. The potential public health impact is substantial - even modest improvements in early detection could significantly reduce GC mortality, addressing a critical healthcare challenge worldwide. The approach is particularly promising for identifying asymptomatic early-stage lesions at a curative stage, potentially bridging the survival gap between countries with and without established screening programs.

