## Supplementary material (Add. file 1, file 2, file 3, Supplementary file) for "Plasma protein biomarkers to detect early gastric preneoplasia and cancer: a prospective study": Supplementary File.docx

**Supplementary Methods**

**Patients diagnosis**

Gastric biopsies from both antrum and corpus were collected during gastric endoscopy. Biopsies were immersed in formalin and processed for haematoxilin-eosin (H&E) staining for histology analysis and diagnosis of gastric lesions. The presence of *H. pylori* was confirmed by Giemsa staining and serology.

### **Plasma mass spectrometry-based proteomics (MS)**

*Plasma proteins depletion and peptidic digestion*

*MARS Hu-14 immunodepletion*

Plasma samples were depleted using the MARS Hu-14 (Agilent, ref: 5188-6560) following the manufacturer's protocol. Briefly, 300µg of total proteins were diluted in Buffer A (part of MARS starter kits of Agilent, ref: 5185-5987), filtered and loaded into the spin column. The non-depleted proteins were then eluted with 2 rounds of 400 µL buffer A by a centrifugation at 100g / 2.5 min / room temperature (RT). Filtrates were combined and further precipitated with Trichloroacetic acid (TCA) 40% (vol:vol) overnight. Samples were washed 2 times with acetone and air-dried before in-solution digestion.

*In-solution digestion*

Depleted samples were resuspended in 100 µL 8M Urea / 100mM NH_4_HCO_3_ denaturation buffer and reduced with 5mM (tris(2-carboxyethyl)phosphine) (TCEP) (Sigma, St Louis, Missouri, USA) for 15 min followed by alkylation with iodoacetamide 20 mM (Sigma, St Louis, Missouri, USA) for 30 min into the dark. Proteins were digested with rLys-C 0.5 µg (Promega, Madison, Wisconsin, USA) for 3h at 37°C and then diluted 9 times for subsequent digestion with Sequencing Grade Modified Trypsin 0.5µg (Promega, Madison, Wisconsin, USA) overnight at 37°C. The digestion was stopped with 4% Formic acid (FA) and peptides were desalted with a reversed phase C18 Stage-Tips method (1). Peptides were then eluted with 80% Acetonitrile (ACN) / 0.1% FA. Finally, samples were dried in vacuum centrifuge and resuspended with 2% ACN / 0.1% FA. For all samples, iRT (Indexed Retention Time) peptides were spiked as recommended by Biognosys.

*Peptide fractionation for spectral library*

A “pool” sample composed of the 40 plasma samples was dedicated to obtaining a spectral library for the data independent acquisition (DIA) approach. The “pool” sample was depleted and digested as indicated above and peptide fractionation was done using poly(styrene-divinylbenzene) reverse phase sulfonate (SDB-RPS) Stage-Tips method as already described (1) (2). Briefly, 3 SDB-RPS Empore discs were stacked on a P200 tip and 7 serial elutions were applied as follow: elution 1 (60mM Ammonium formate (AmF) / 20% ACN / 0.5% FA), elution 2 (80mM AmF / 30% ACN / 0.5% FA), elution 3 (95mM AmF / 40% ACN / 0.5% FA) , elution 4 (110mM AmF / 50% ACN / 0.5% FA), elution 5 (130mM AmF / 60% ACN / 0.5% FA), elution 6 (150mM AmF / 70% ACN / 0.5% FA) and elution 7 (80% ACN / 5 % ammonium hydroxide). All fractions were dried and resuspended in 2% ACN / 0.1% FA before injection. For all fractions, iRT peptides were spiked as recommended by Biognosys.

*Peptide elution and mass spectra acquisition*

For each sample, 1 μg of peptides was injected onto a 44cm (DDA acquisitions) or 50cm (DIA acquisitions) home-made C18 column (1.9 μm particles, 100 Å pore size, ReproSil-Pur Basic C18 - Dr. Maisch GmbH, Ammerbuch-Entringen, Germany) after an equilibration step in 100 % solvent A (H_2_O, 0.1% FA). Peptides were eluted with a multi-step gradient from 2 to 7 % solvent B (80 % ACN, 0.1 % FA) during 5 min, 7 to 23 % solvent B during 70 min, 23 to 45 % solvent B during 30 min and 45 to 95 % solvent B during 5 min at a flow rate of 250 nL/min over 132 min. Column temperature was set to 60°C.

*Data Dependent Acquisitions (DDA) for spectral library generation*

Mass spectra were acquired using Xcalibur software using a data-dependent Top 10 method with a survey scans (300-1700 m/z) at a resolution of 60,000 and a MS/MS scans (fixed first mass 100 m/z) at a resolution of 15,000. The automatic gain control (AGC) target and maximum injection time for the survey scans and the MS/MS scans were set to 3.0E+06, 100ms and 1.0E+05, 45ms respectively. The isolation window was set to 1.6 m/z and normalized collision energy fixed to 28 for high-energy collisional dissociation (HCD) fragmentation. We used a minimum AGC target of 2.0E+03 for an intensity threshold of 4.4E+04. Unassigned precursor ion charge states as well as 1, 7, 8 and >8 charged states were rejected and peptide match was disable. Exclude isotopes was enabled and selected ions were dynamically excluded for 45 seconds.

*Data Independent Acquisitions (DIA) from plasma samples*: mass spectra were acquired under data-independent acquisition mode with the XCalibur software. Each cycle was built up as follows: one full MS scan at resolution 60 000 (scan range between 349 and 1214 m/z), AGC was set at 3.0E+06 and maximum injection time was set at 60 ms. All MS1 was followed by 36 isolation windows of 25 m/z, covering the MS1 range. The AGC target was 2.0E+05 with an automatic maximum injection time and normalized collision energy (NCE) was set to 28. All acquisitions were done in positive and profile mode.

***Data processing for protein identification and quantification***

*Building of the spectral library:* Raw data were analyzed using MaxQuant software version 1.5.0.30 (3) using the Andromeda search engine (4). The MS/MS spectra were searched against the Human SwissProt database (20,203 entries the 12/04/2018). Variable modifications (methionine oxidation and N-terminal acetylation) and fixed modification (cysteine carbamidomethylation) were set for the search and trypsin with a maximum of two missed cleavages was chosen for searching. The minimum peptide length was set to 7 amino acids and the false discovery rate (FDR) for peptide and protein identification was set to 0.01. The main search peptide tolerance was set to 4.5 ppm and to 20 ppm for the MS/MS match tolerance. Second peptides were enabled to identify co-fragmentation events.

*Data analysis for DIA method*: DIA experiments were analyzed using Spectronaut X (v. 13.2.190705.43655 Biognosys AG). Dynamic mass tolerance at the MS1 and MS2 levels was employed. The XIC RT Extraction Window was set to dynamic with a correction factor of 1. Calibration mode was set to automatic with nonlinear iRT calibration and precision iRT enabled. Decoys were generated using the mutated method and a dynamic limit. P-value estimation was performed using a kernel density estimator. Interference correction was enabled with no proteotypicity filter. Major grouping was by Protein-Group ID, and minor grouping was by stripped sequence. The major group quantity was mean peptide quantity. The major group top N was enabled with a minimum of 1 and a maximum of 3. Minor group quantity was mean precursor quantity. The minor group top N was enabled with a minimum of 1 and a maximum of 3. The quantity MSLevel was MS2, and quantity type was area. Q value was used for data filtering. Cross run normalization was enabled with Q value sparse for the row selection and local normalization for the strategy. The default labeling type was label-free with no profiling strategy and unify peptide peaks not enabled. The protein inference workflow was set to automatic.

**Legend Supplementary Figures:**

**Figure S1:** **Volcano plots corresponding to the “relaxed” differential analysis of LC-MS/MS data.** The “relaxed” analysis is performed by imposing at least 2 quantified values among the patients of one of the two compared categories. For complete lists of differentially abundant proteins, see additional file 1.

**Figure S2: Volcano plots corresponding to the “strict” differential analysis of LC-MS/MS data.** The “strict” analysis is performed by imposing at least 9 quantified values among the patients of one of the two compared categories. For complete lists of differentially abundant proteins, see additional file 3.

**Figure S3: Plasma concentration level of the 15 protein candidates identified by LC-MS/MS analysis according to gender**. Protein levels were measured by ELISA commercial assays as indicated in the material and methods section, on all samples of the cohort. F: female (light color), M: male (dark color). Statistical analysis was performed using Mann-Whithey test**.** Only significant differential levels are indicated **(******P<0.0001; ***P<0.001; **P<0.01; *P<0.05)

**Figure S4:** **Plasma concentration level of the 15 protein candidates identified by LC-MS/MS analysis according to age**. Protein levels were measured by ELISA commercial assays as indicated in the material and methods section, on all samples of the cohort, and compared in patient groups and healthy controls younger or older than 50 years old. <50: under 50 years-old (light color); ≥50: up 50 years-old (dark color). Data corresponding to the 2 patients under 50 years-old with preneoplasia are represented by scatter dot plots. Statistical analysis was performed using Mann-Whithey test**.** Only significant differential levels are indicated **(******P<0.0001; ***P<0.001; **P<0.01; *P<0.05).

| **Protein name** | **Accession number** | **Reference ELISA assays** |
| --- | --- | --- |
| ARG-1 | P05089 | MBS2509985 |
| ATAD3B | Q5T9A4 | MBS7241561 |
| CA2 | P00918 | MBS705181 |
| DCD | P81605 | MBS062371 |
| F13A1 | P00488 | MBS702048 |
| HPT | P00738 | DY8465 |
| IGFALS | P35858 | MBS2020386 |
| JUP | P14923 | MBS2018947 |
| KIF20B | Q96Q89 | MBS9321702 |
| KRT14 | P02533 | MBS4500642 |
| KRT19 | P08727 | MBS2503663 |
| LBP | P18428 | MBS4500696 |
| LEP | P41159 | DY398 |
| MAN2A1 | Q16706 | MBS7212732 |
| S100A12 | P80511 | MBS773252 |

**Table S1: List of protein biomarker candidates identified by mass spectrometry and quantified by ELISA on all the samples of the cohort.** MBS and DY references are from MyBiosource, USA and R&D systems, USA, respectively.

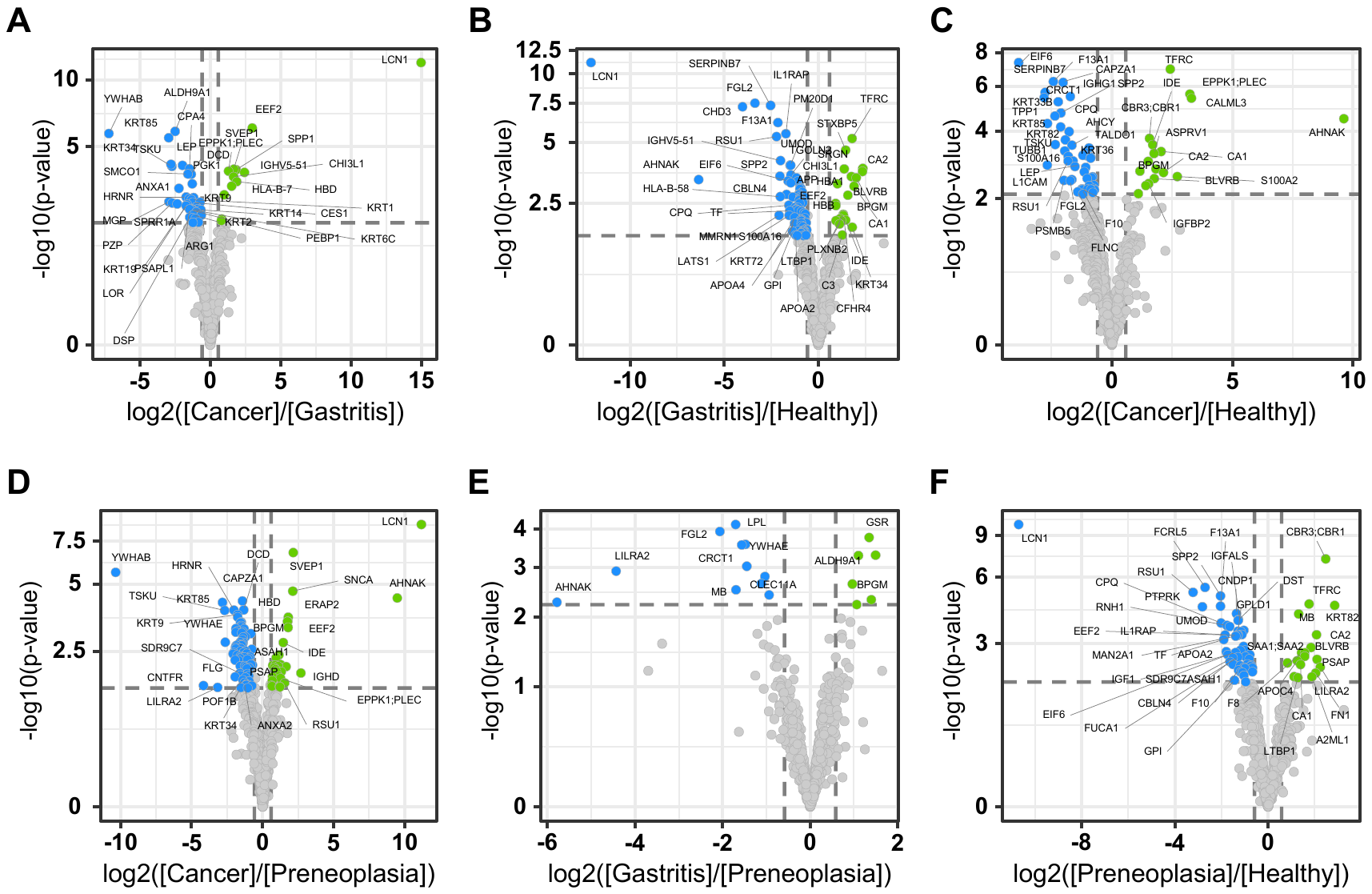
**Figure S1**

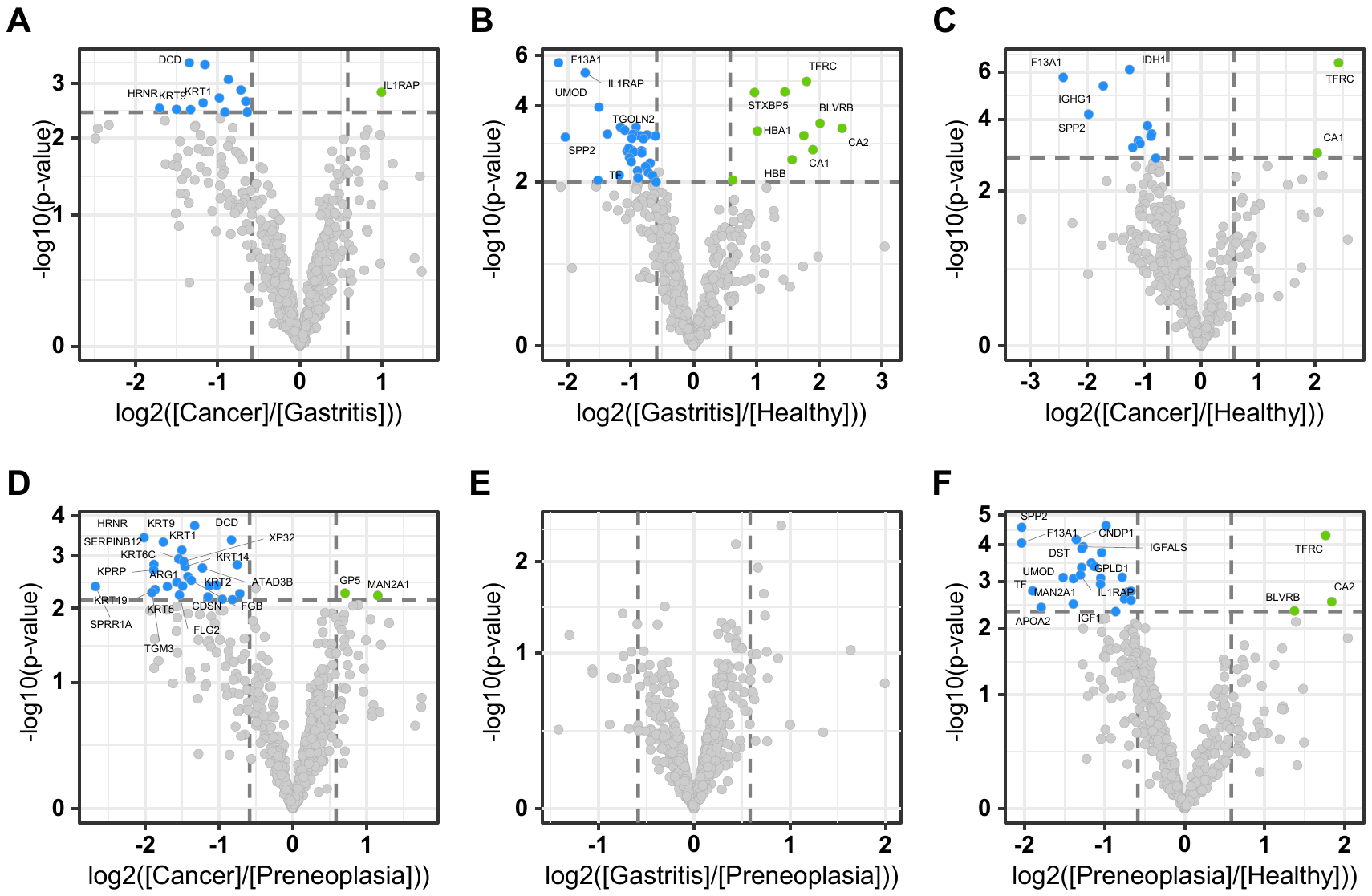
**Figure S2**

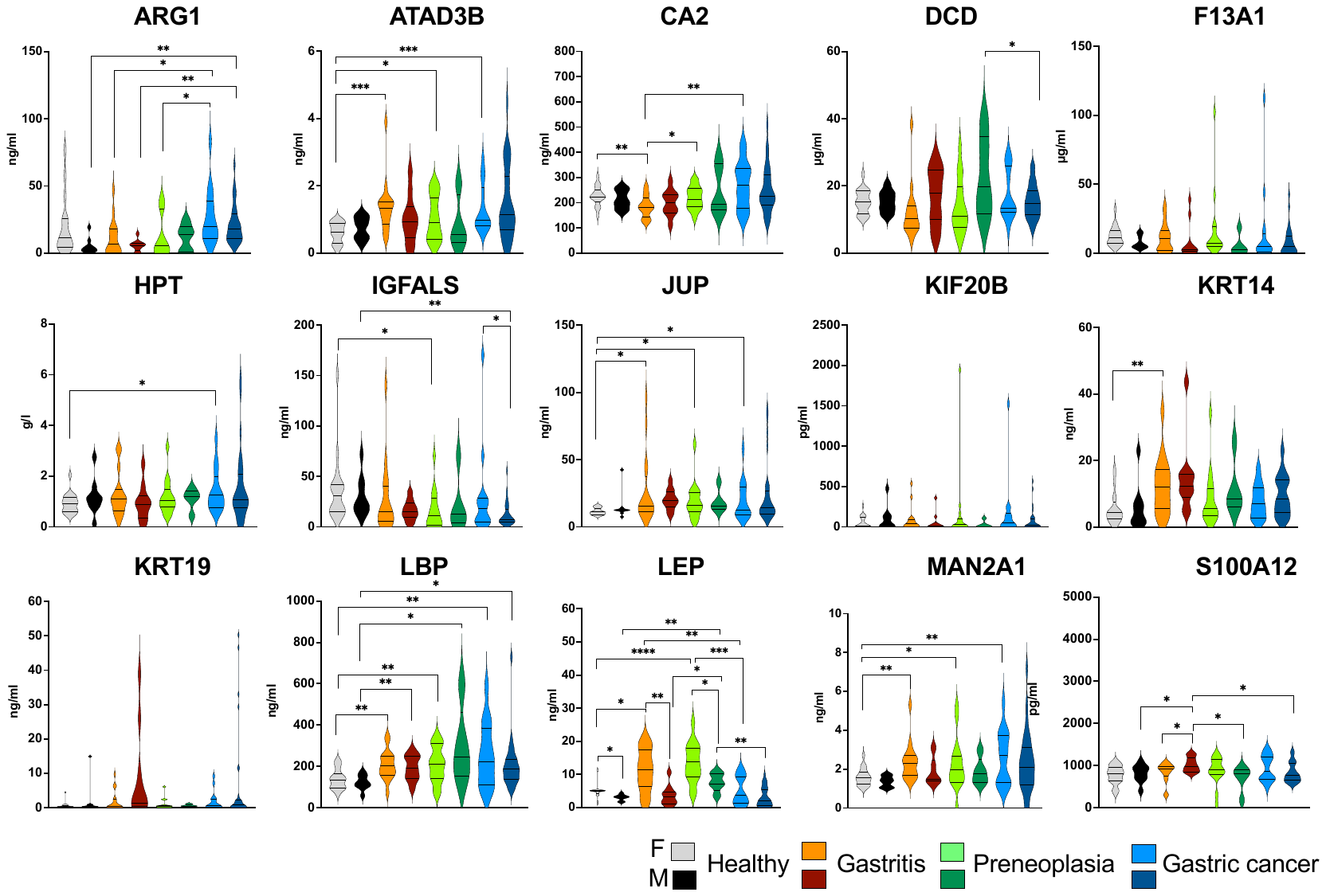
**Figure S3**

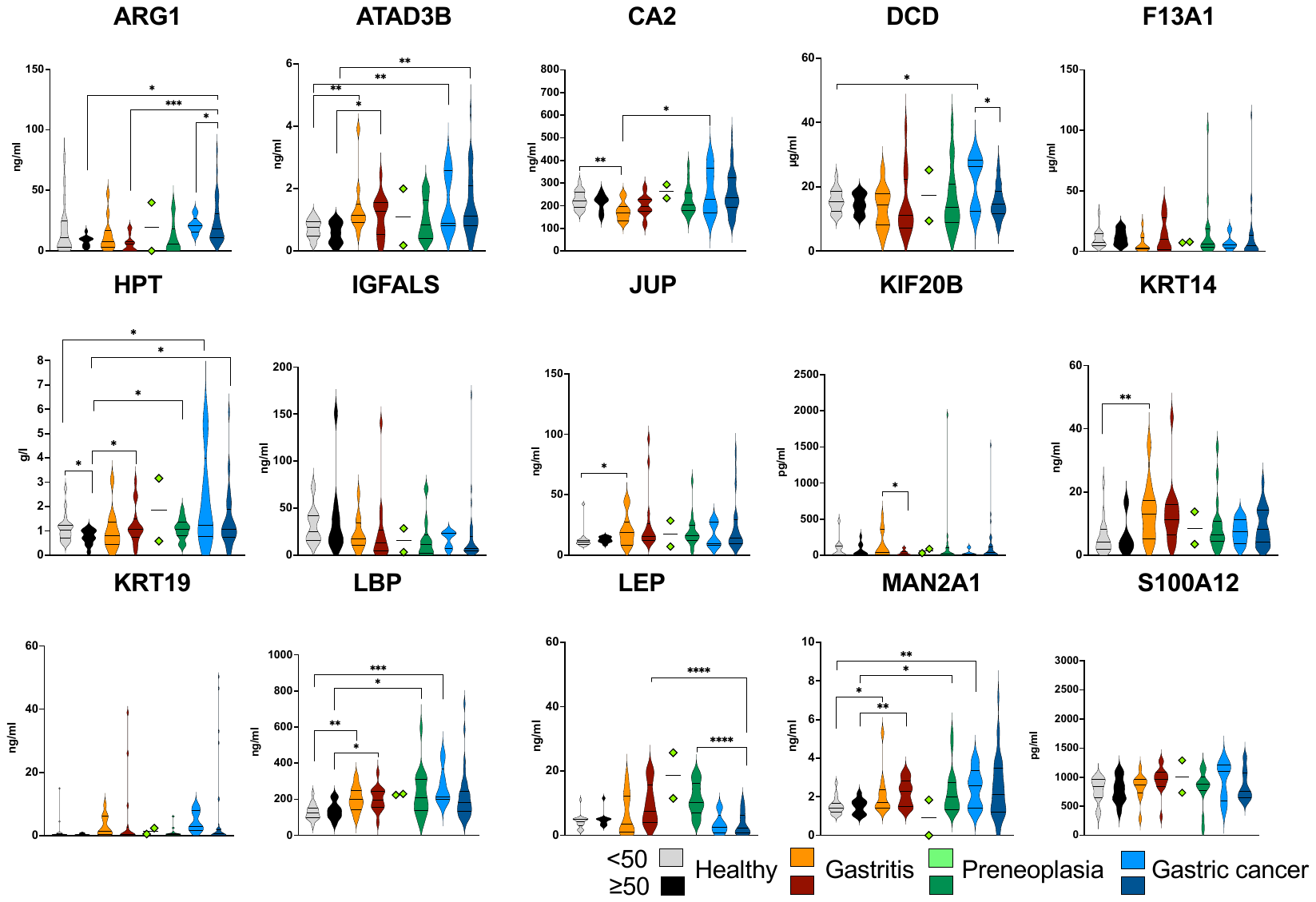
**Figure S4**

**Additional files**

**Additional file 1 (xlsx file):** Excel file containing the statistical results of the “relaxed” analysis

**Additional file 2 (xlsx file):** Excel file containing the statistical results of the “strict” analysis

**Additional file 3 (xlsx file):** The means and standard deviations of the Area Under the Receiver Operating Characteristic Curve (AUROC) values obtained through 4-fold cross validation, repeated 10 times from the ELISA data of the 138 patients in the cohort for all tested combinations of biomarkers.
